# Social modulation of activity and orientation fosters complex interactions in young zebrafish

**DOI:** 10.64898/2026.09.04.749288

**Authors:** Laura Desban, Karen Guillemin, Judith S. Eisen, Raghuveer Parthasarathy

**Affiliations:** Institute of Neuroscience and Department of Biology, University of Oregon, USA; Institute of Molecular Biology and Department of Biology, University of Oregon, USA; Department of Physics, University of Oregon, USA

**Keywords:** Social behavior, proximity behavior, physical contacts, turning behavior, zebrafish

## Abstract

Zebrafish are an important model organism for investigating collective behavior and neuropsychiatric disorders due to attributes such as fast development, high genetic homology with humans and a rich behavioral repertoire. Social behavior in zebrafish emerges as early as twelve days post-fertilization, building toward diverse, coordinated, and complex adult interactions. Most studies on social phenotypes, including in zebrafish, have focused on adult animals and assess isolated traits such as aggression, boldness, or social memory, leaving the critical period of early development when sociality emerges and the mechanisms underlying social behavior establishment largely unexplored. We present here a behavior analysis pipeline that identifies features describing general locomotion and social behavior based on simple geometric criteria. We apply this analysis to pairs of freely interacting two-week-old zebrafish and characterize nascent social preference. We find that young zebrafish favor a specific range of inter-fish distances, creating a social space characterized by specific modulation of locomotion features. Through quantitative analysis and modeling, we show that this proximity behavior is largely established through a turning bias toward counterparts and a socially induced suppression of the variance in turning angles over broader ranges of inter-fish distances. Furthermore, we describe how proximity fosters engagement by young zebrafish in transient, complex short-range physical interactions that include various forms of inter-fish contacts, reminiscent of patterns described in adult animals. Our approach has broad applications for understanding the ontogeny of sociality and collective behaviors and describing subtle phenotypes associated with various human disorders.

**Graphical abstract:** 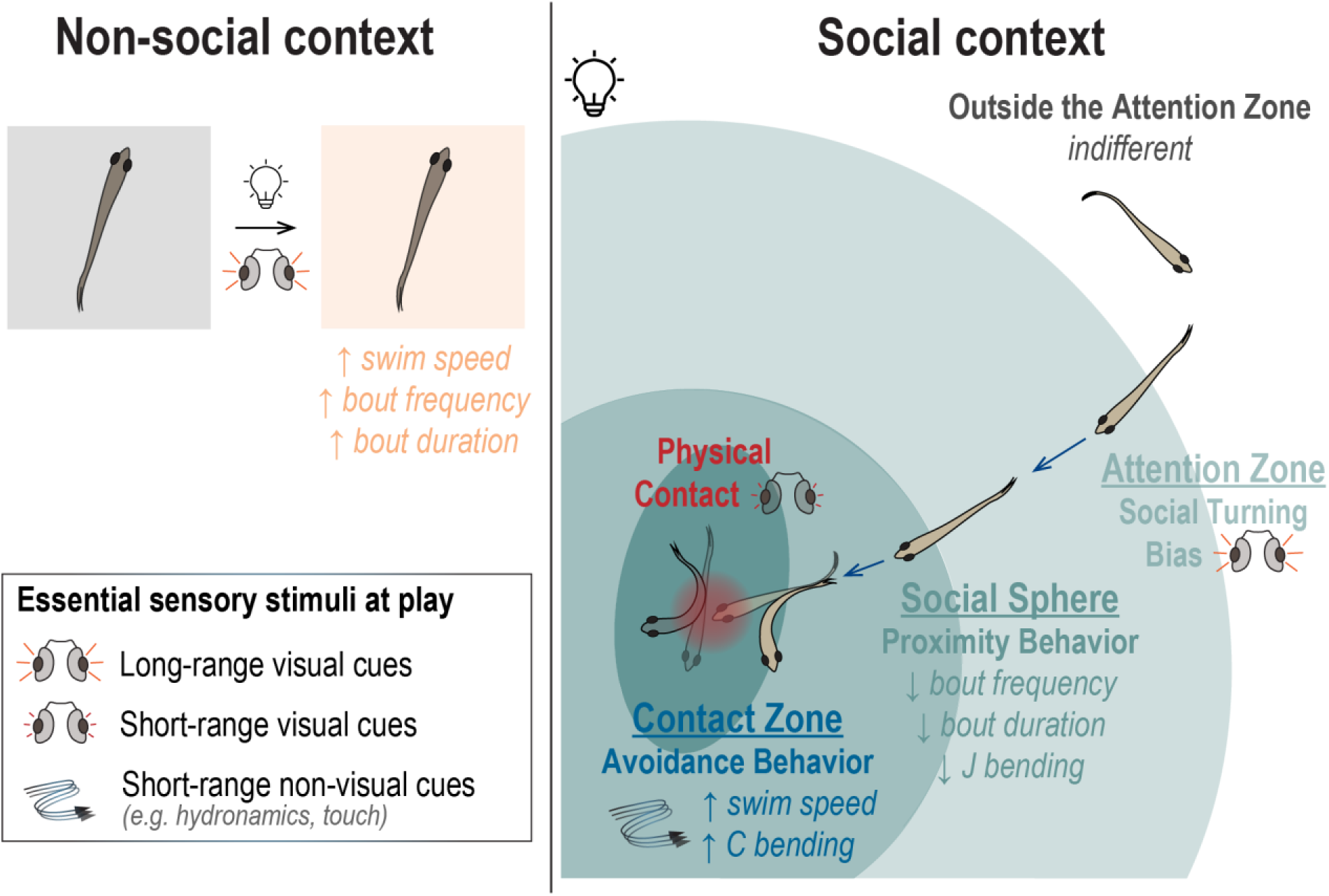

## Introduction

Interactions between individuals are critical for the development and well-being of social animal species, contributing to fundamental functions such as reproduction, feeding, avoidance of threats, and response to stress (Heinrichs et al., 2003; Holt-Lunstad et al., 2015; Ward & Webster, 2016). In humans, deficits in sociality are associated with higher prevalence of neuropsychiatric and neurodevelopmental conditions such as autism spectrum disorder, post-traumatic stress disorder, depression, and schizophrenia (Frye, 2018; Porcelli et al., 2019). The incidence of these conditions is increasing and, while a growing body of evidence points toward a complex interplay between multiple genetic and environmental factors, their etiology and pathogenesis remain unclear. Animal models are critical to enable studies that illuminate the neural mechanisms underlying complex behavioral traits and disorders(Bale et al., 2019; Nestler & Hyman, 2010). Over the past decade, zebrafish have proved to be a useful model system, thanks to sharing over 70% genetic homology with mammals including humans, conserved neurodevelopment and neural structures, and complex sociality (Apukhtin et al., 2026; Fontana et al., 2018; Sakai et al., 2018). Zebrafish’s high fecundity, short generation time, external development and genetic tractability are additional features that make it particularly amenable to large-scale screens and genetic manipulations to investigate neurodevelopmental mechanisms underlying behavioral disorders.

Zebrafish are social animals that engage in diverse, complex social interactions (Buske & Gerlai, 2011; Saverino & Gerlai, 2008). Social behavior in zebrafish emerges as early as 12 days post-fertilization (dpf) in the form of social preference largely mediated by visual cues (Dreosti et al., 2015; Engeszer et al., 2004; Nunes et al., 2020; Stednitz & Washbourne, 2020). As zebrafish mature, their social behaviors rapidly develop to give rise to complex group behaviors such as shoaling and schooling. Currently, most investigations of social behavior and associated deficits focus on adults and target specific traits in isolation such as shoal cohesion, social memory, or aggression (Ogi et al., 2021; Pham et al., 2012). While informative, this trait-by-trait approach fragments sociality into discrete components and lacks essential contextualization to study social behavior as a whole in freely interacting animals. Furthermore, the focus on adults has left early developmental stages understudied, and the extensive and critical changes that occur at younger stages and will eventually lead to normal versus aberrant adult neurobehavioral phenotypes poorly mapped.

Characterization and quantification of animal behavior is an enduring challenge that has spurred a variety of strategies. Recent studies have applied machine learning methods and unsupervised approaches to extract biologically relevant, recurrent structures in recordings of freely behaving zebrafish (Johnson et al., 2020; O’Shaughnessy et al., 2024; Reddy et al., 2022; Stednitz et al., 2025). However, identifying robust or interpretable behavioral patterns has proven difficult. While automatically extracted motifs can potentially be explained in human-comprehensible terms, as shown recently for dominance contests in adult zebrafish (O’Shaughnessy et al., 2024), such interpretation is generally challenging. In addition, the applicability of machine-learned motifs to data collected across different laboratories and different experimental conditions remains unclear. In contrast, the classic approach to ethology of observation and description provides interpretable insight but is labor intensive and prone to biases. Here, we undertake a hybrid approach that combines human-identified behavioral patterns with quantitative analysis and automated identification, applied to image-derived tracking data from recordings of freely interacting pairs of two-week-old zebrafish. We construct a minimal model of zebrafish navigation yielding predictions for characteristics of pair-wise social interactions across relevant spatial and temporal scales that can be compared to human observations. In our model, rapid, short-range interactions between pairs of fish are identified from time-series data without segmentation into bouts of swimming, allowing recognition of patterns independent of locomotion structure.

Our results uncover previously undescribed features of social behavior, revealing that social context strongly impacts locomotion and how individuals navigate their environment. Our analysis indicates that young zebrafish actively maintain a preferred range of inter-individual distances by biasing turns toward conspecifics and reducing turning noise. The latter mechanism, a reduction in stochastic movements that would be interpreted as increased attention or awareness of a proximal social partner, appears to be the dominant factor in the maintenance of proximity. Young zebrafish also engage in brief physical contacts with their partner. These transient interactions have not previously been reported at this developmental stage and are reminiscent of contact-based behaviors described in adult social repertoires (Kalueff et al., 2013). All together, these findings reveal that two-week-old zebrafish are active social agents that implement structured strategies to navigate social space. Our observations establish a quantitative foundation for dissecting the environmental, sensory, neural and genetic mechanisms underlying social behavior and characterizing subtle phenotypes present at the onset of sociality that could develop into social deficits at later stages (Petersen et al., 2026).

## Results

### Social context impacts two-week-old zebrafish overall locomotion

To characterize social behavior of two-week-old zebrafish, we recorded freely swimming individuals placed in shallow circular arenas for ten minutes (Fig. 1A). Animals were recorded either alone, to assess baseline locomotion, or in pairs of size-matched individuals to assess social behavior. Since visual stimuli are known to be instrumental in zebrafish sociality (Dreosti et al., 2015; Engeszer et al., 2004, 2007; Nunes et al., 2020; Stednitz & Washbourne, 2020), we performed half the recordings under homogeneous illumination (30 individual fish, 29 pairs) and half in the dark (29 individual fish, 30 pairs), allowing us to distinguish between social behavior components that are visually driven or not, respectively (Fig. 1B). We then extracted behavioral features using a newly established automated analysis pipeline that uses body position tracking data to calculate individual locomotion and pairwise social parameters in each frame (Fig. 1C-F; see Methods).

**Figure 1.**
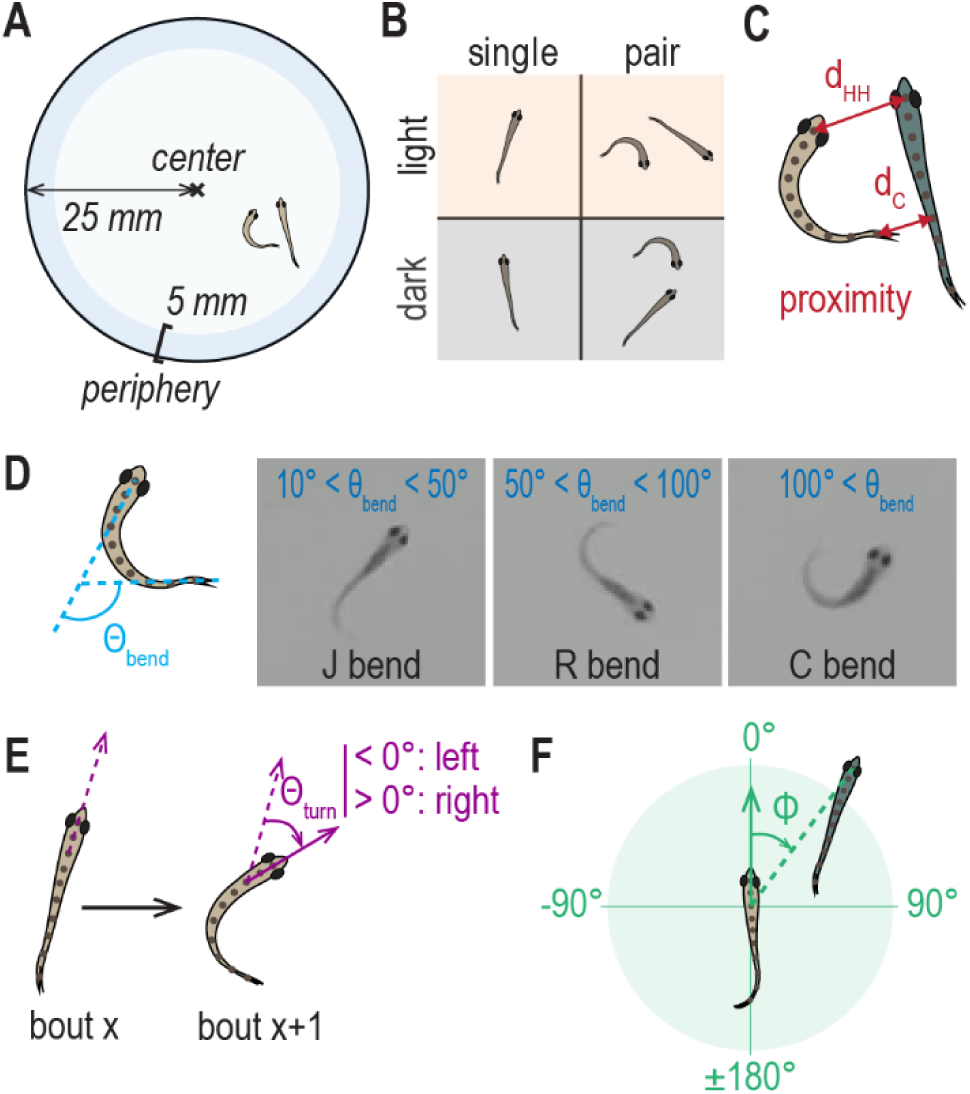
Analysis of locomotion and social behavior features based on geometric criteria. **A**, Schematic of the circular behavioral chambers used to record freely swimming two-week-old zebrafish. **B**, Our dataset consists of recordings of single individuals and pairs of two-week-old zebrafish, in dark or light conditions. Several behavioral features extracted by our pipeline are illustrated here, including measures of inter-fish distance (**C**), either head-head distance (d_HH_) or closest distance (d_C_), tail bending angle (Ɵ_bend_, **D**) with subcategorization of bends as J bends, R bends or C bends, bout-to-bout turning angle (Ɵ_turn_, **E**), and relative orientation angle (ф, **F**) quantifying the relative position between the two fish from the focal fish standpoint.

We first analyzed the behavior of single individuals to establish baseline locomotion and compared between dark and light conditions to identify features dependent on illumination (Fig. 2, Table 1). Overall, we observed robust light-induced activation of locomotion in two-week-old zebrafish. We found that single individuals spend a greater fraction of time swimming in lit arenas than in the dark (Fig. 2A), reflected both in longer swim bout durations (Fig. 2B) and shorter inter-bout intervals (Fig. 2C), and swim on average faster (Fig. 2D). Conversely, other features associated more with how two-week-old zebrafish occupy and explore their environment, such as turning speed (Fig. 2E), frequency of turning behavior (Fig. 2F,H,I), and time spent at the periphery (Fig. 2G) did not vary between dark and light conditions.

**Figure 2.**
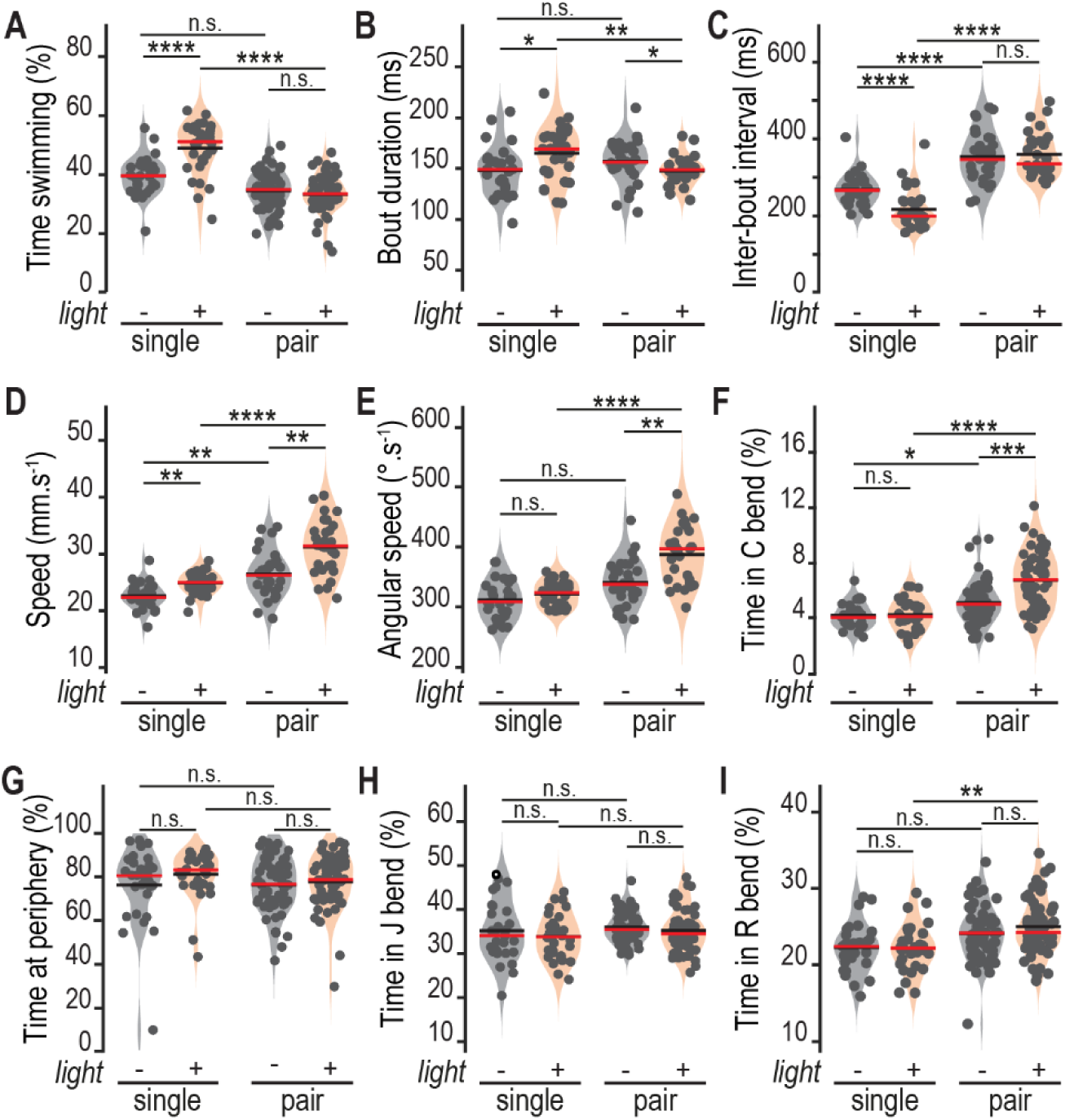
Social context impacts two-week-old zebrafish overall locomotion. Social context impacts aspects of locomotion in three different ways: (i) overall damping of light-induced increase of activity otherwise observed in single individuals (**A-C**) with reduced time spent swimming (**A**) due to concomitant decrease in average bout duration (**B**) and increase in inter-bout interval (IBI, **C**); (ii) synergistic effect with lighting (**D-F**) observed as increase in average speed when moving (**D**), average angular speed when moving (**E**) and average time spent performing C bends (**F**); and (iii) no effect (**G-I)** on average time spent at the periphery (**G**), and performing J bends (**H**) or R bends (**I**). Each individual dot represents a single fish; for pairs, each dot represents a single fish within a pair (A,F-I) or the average between two fish in a pair (B-E). Red and black bars in graphs indicate the median and the mean, respectively. Statistical significance was tested using two-sample Kolmogorov-Smirnov tests and corrected using the Bonferroni method for multiple comparisons. Median values and standard deviations for each measure are reported in **Table 1**.

**Table 1.** Social context impacts two-week-old zebrafish locomotion. Medians and standard deviations across groups for all parameters considered in Fig. 2, categorized depending on how they are affected by social context: (i) damping of light-induced increase in activity (orange box), (ii) synergistic effect with lighting (red box) and (iii) no effect (grey box).

| | | Swimming<br>% | Bout<br>duration<br>ms | Inter-bout<br>interval<br>ms | Speed<br>$mm.s^{-1}$ | Angular<br>speed<br>$^{\circ}.s^{-1}$ | C bend<br>% | Periphery<br>% | J bend<br>% | R bend<br>% |
| --- | --- | --- | --- | --- | --- | --- | --- | --- | --- | --- |
| single | dark | 40 ± 6.6 | 149 ± 24 | 266 ± 42 | 22 ± 2.3 | 309 ± 32 | 4.1 ± 0.9 | 81 ± 18 | 34 ± 6.8 | 22 ± 3.1 |
|  | light | 51 ± 8.8 | 169 ± 26 | 199 ± 51 | 25 ± 1.9 | 324 ± 21 | 4.2 ± 1.1 | 83 ± 11 | 34 ± 5.2 | 22 ± 3.1 |
| pair | dark | 35 ± 6.5 | 156 ± 21 | 346 ± 67 | 26 ± 4.1 | 338 ± 38 | 5.1 ± 1.5 | 77 ± 12 | 35 ± 3.5 | 24 ± 3.5 |
|  | light | 33 ± 6.3 | 149 ± 14 | 335 ± 57 | 31 ± 4.8 | 397 ± 47 | 6.8 ± 2.0 | 79 ± 12 | 35 ± 4.9 | 24 ± 3.7 |

We then looked at the behavior of pairs of fish to see how sociality impacted general locomotion (Fig. 2; Table 1). We observed that the presence of another conspecific in the arena dampens light-induced increase in swimming activity described above for single individuals, and significantly reduces frequency of swim bouts in a light-independent fashion (Fig. 2A-C; Table 1, orange box). Conversely, we found a synergistic effect of sociality with light (Fig. 2D-F; Table 1, red box) on swimming speed, swimming angular speed and fraction of time spent in C bends. C bends are typically described as high angular speed tail bending events in the context of escape response (Budick & O’Malley, 2000a; Mirat et al., 2013), indicating that avoidance behavior may take place in a social context when another individual is present. Finally, the presence of another conspecific did not affect how much time zebrafish spend at the periphery, or in J and R tail bends (Fig. 2G-I; Table 1, grey box), two types of ‘routine’ turning behaviors displayed by zebrafish to reorient during navigation (Budick & O’Malley, 2000a; Marques et al., 2018; Mirat et al., 2013).

Together, these observations indicate that two-week-old zebrafish significantly modulate their locomotion, but not how they occupy space, depending on lighting condition and social context. Notably, in the presence of another individual in the arena, they swim less frequently but faster, and exhibit movements reminiscent of rapid avoidance.

### Two-week-old zebrafish actively maintain a preferred range of social distance through social turning bias

Our locomotion analysis did not reveal gross effects of lighting or social context on how two-week-old zebrafish occupy space in behavioral arenas. Overall, zebrafish in all tested groups spend approximately 80% of their time at the periphery (Fig. 2G), exhibiting previously described thigmotaxis behavior (Champagne et al., 2010; Schnörr et al., 2012). Nonetheless, we reasoned that paired zebrafish could still modulate space occupation relative to one another. We explored this hypothesis by analyzing various measures of proximity between two individuals (Fig. 1C). Given the robust thigmotaxis behavior noted above, the null expectation for the proximity distribution of non-interacting pairs of fish is not uniform, but instead a nontrivial function of the arena geometry and the radial distribution of fish positions (Supp. Fig. 4,6). Therefore, to generate control distributions for comparisons, we cyclically shifted in time the positions of one fish in each pair by half the recording duration, generating time-shifted datasets that retained fish locomotion characteristics. Values derived from these control datasets were used throughout the analysis of social behavior, displayed in graphs with dotted lines as opposed to solid lines for true datasets.

First, we measured inter-fish distance as the distance between the heads of the two fish (head-head distance or d_HH_; Fig. 1C). We extracted this measure at each frame and calculated the average across the entire recording for each pair (Fig. 3A) and found that pairs under illumination swim significantly closer (median d_HH_ = 21 mm ± 5 mm in the light versus 26 mm ± 5 mm in the dark, and 25 mm ± 4 mm in control time-shifted light data). We further compared the probability density distributions, i.e., the distributions of the likelihood for each distance value of each group, as well as control distributions for both light and dark conditions. Head-head distance distributions revealed that, under illumination, paired two-week-old zebrafish strongly favor distances in the 5-15 mm range (Fig. 3B) while pairs in the dark show an opposite trend compared to control datasets, suggesting that maintaining or avoiding short inter-fish distances in light and dark settings, respectively, are active processes. These observations are consistent with reports in the literature that zebrafish exhibit robust preference for shorter inter-fish distances from 12 dpf onward (Stednitz & Washbourne, 2020) and that social behavior is strongly visually driven at this stage (Dreosti et al., 2015; Engeszer et al., 2004, 2007).

**Figure 3.**
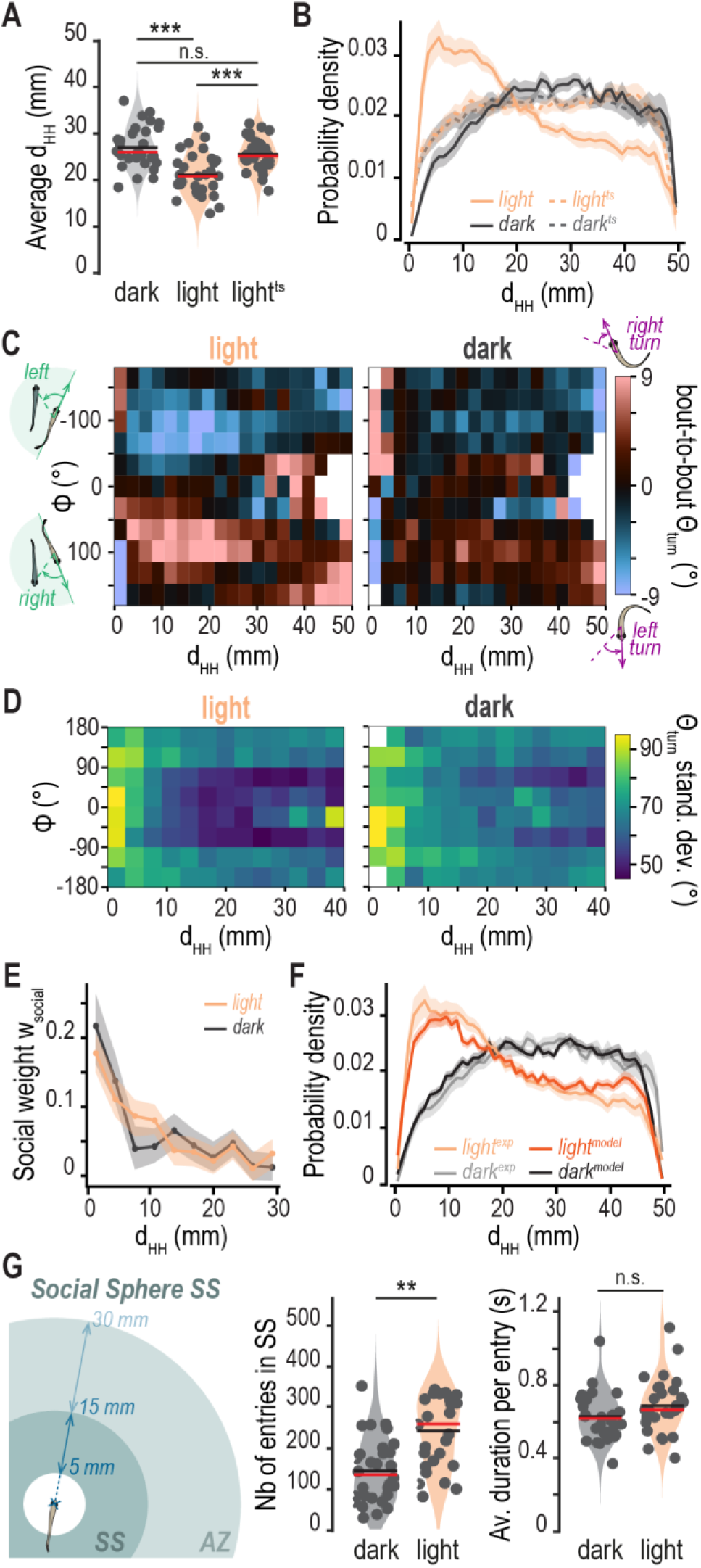
Two-week-old zebrafish actively maintain a preferred range of social distance through social turning bias. In lit arenas, the average head-head distance (d_HH_) between two fish is significantly reduced compared to that in the dark (**A**) and the overall probability distribution is skewed toward short inter-fish distances (**B,** dashed lines indicate control time-shifted datasets light^ts^ and dark^ts^). **C**, Two-dimensional plots of the average bout-to-bout turning angle (Ɵ_turn_) as a function of d_HH_ and relative orientation (ф). In lit arenas, two-week-old zebrafish exhibit a turning bias toward the other fish for d_HH_ in the 5-30 mm range. Conversely, they turn away from each other at shorter distances (< 5 mm) and this effect is stronger in the dark. **D**, Two-dimensional plots of the standard deviation of the bout-to-bout turning angle as a function of d_HH_ and ф. The variation of Ɵ_turn_ is high at short distances (< 5 mm) but especially low for d_HH_ in the range of 5-30 mm in lit arenas. **E**, The genuine social weight w_social_ decays to zero by about 20 mm in lit arenas versus 7 mm in dark arenas. **F**, Comparison of the probability distribution of d_HH_ from experimental data (light^exp^ and dark^exp^) and from the simulation (light^model^ and dark^model^) for the full model as detailed in Supp. Text 1. **G**, The Attention Zone (AZ) is the zone where zebrafish start showing social bias of turning behavior and the Social Sphere (SS) is the preferred range of d_HH_ (left schematics). Pairs in the light enter the social sphere significantly more often (middle graph) but stay on average for the same duration as pairs in the dark (right graph). Each individual dot represents a pair. Traces in B, E and F represent the average across all pairs per condition, and the shaded area indicates the s.e.m. White squares in C and D indicate bins with no data. Red and black bars in graphs indicate the median and the mean, respectively. Statistical significance was tested using two-sample Kolmogorov-Smirnov tests.

To better understand how two-week-old zebrafish establish and maintain this preferred inter-fish distance range, we asked whether a minimal behavioral model could reproduce the observed d_HH_ distributions. We hypothesized a model in which decisions are instantaneous, i.e., without memory (Markovian), but in which bout behavior is influenced by position and orientation relative to the other fish and to the arena wall. We first analyzed and modeled the behavior of single fish. As described in detail in Supplementary Text 1 and associated supplementary figures, modeling fish trajectories as random walks, with steps drawn from the observed set of bout displacements binned by distance from the arena center (r) and alignment angle with the wall (ψ) reproduces the radial distance distributions observed in experimental data (Supp. Fig. 4). Importantly, neglecting the wall alignment angle leads to poor agreement with the data, highlighting the prominence of thigmotaxis behavior.

We next modeled pair behavior, again based on a Markovian random walk for each fish. We studied the data to determine the key drivers determining bout behavior in relation to position relative to the other fish. We found that change in turning angle between bouts is strongly biased depending on social parameters, namely the head-head distance (d_HH_) and relative orientation (φ) that quantifies the angle between the forward direction of the focal fish and the location of the other fish’s head (Fig. 1F). For distances between 5 and 30 mm, fish are more likely to turn to the left (θ_T_ < 0) if the other fish is to the left (φ < 0), and to the right (θ_T_ > 0) if the other fish is to the right (φ > 0) (Fig. 3C). This is in part a geometric constraint: if a wall is to the right, neither the other fish nor a turn can be to the right, resulting in a residual environmental bias of turning behavior as can be observed in control time-shifted datasets (Supp. Fig. 1). However, this bias was essentially absent in dark arenas (Fig. 3C), suggesting a real social component to turning that is visually driven, consistent with previous reports. Furthermore, we also observed the opposite tendency at shorter distances (i.e., under 5 mm) when fish turn away from each other (Fig. 3C). This behavior occurring at short distances was much stronger in pairs in the dark than in pairs with access to visual cues, and suggested the triggering of avoidance responses upon close proximity, especially in situations when zebrafish cannot see each other.

Motivated by the evidence of socially biased turning behavior, we tested the hypothesis that instantaneous turning preferences suffice to explain the observed preferred inter-fish distancing. Our minimal model begins with bout lengths and durations sampled from the data, binned by radial position (r) and wall alignment angle (ψ) as in the model for single individuals. The turning angle at each simulated bout is composed of two components, an ‘intrinsic’ turning angle sampled from single fish data and a ‘social’ turning angle given by the inter-fish geometry, the latter being simply the relative orientation of the other fish, φ (Fig. 1F). The combination of these angles involves a weighted sum and a stochastic factor, both of which are derived from the data and depend on inter-fish distance (Suppl. Text 1). We find that the ‘social weight’, i.e., the fraction of the overall contribution to the turning angle from the inter-fish orientation, decays with inter-fish distance, reaching zero by about d_HH_ = 15 mm in the light and 7 mm in the dark (Fig. 3E; Supp. Text 1 and Supp. Fig. 8), implying integration of different sensory modalities at different inter-fish distances, namely visual cues versus non-visual cues at long versus short range, respectively. The stochastic factor is motivated by a surprising observation: the standard deviation of turning angles in each (r, ψ) bin shows a strong dependence on inter-fish distance and relative orientation (Fig. 3D; Supp. Text 1 and Supp. Fig. 9). For fish in lit arenas, the variance is low if the other fish is roughly 10-30 mm away and forward (|φ| < 90°), a pattern absent in the dark. At very close separations (d_HH_ < 5 mm), the variance is large independent of lighting condition. Rescaling the distribution of sampled, weighted turning angles by the ratio of the observed standard deviation to its asymptotic, long-distance value accounts for both the average social turning bias and the socially mediated variance (Supp. Text 1). With these modulated parameters in hand, we performed simulations of pairs of fish under this model and compared the resulting distributions of radial positions (Supp. Fig. 6) and head-head distances (Fig. 3B) to experimental data. The agreement is remarkably good for both light and dark conditions. We emphasize that the model requires zero adjustable parameters and is, in contrast to other recent models of zebrafish and fish behavior (Groneberg et al., 2020; Harpaz et al., 2017, 2026; Johnson et al., 2020; Marques et al., 2018; Stednitz et al., 2025), memoryless, capturing key aspects of the data without invoking persistent behavioral states. Our analysis implies that two-week-old zebrafish establish and maintain preferred distancing essentially through the modulation of turning behavior, the largest factor being socially structured turning variance. As discussed below, this control of variance may be a previously unrealized mechanism of behavioral modulation in animal contexts.

These observations allowed us to define an ‘attention zone’ (AZ, d_HH_ < 30 mm) where zebrafish, using long-range visual cues, exhibit less noisy and socially biased turning behavior, thereby establishing the ‘social sphere’ (SS), i.e., the actively maintained, preferred range of head-head distances (5-15 mm) (Fig. 3G, left schematic). We then quantified how often and for how long zebrafish enter the social sphere. Pairs under illumination enter this space significantly more often than pairs in the dark (Fig. 3G, middle panel; 260 ± 85 entries in the light versus 136 ± 79 entries in the dark) but spend the same amount of time there (Fig. 3G, right panel; median duration = 660 ms ± 154 ms in the light versus 612 ms ± 129 ms in the dark), suggesting that visual cues are important for initiating entry, but not for staying in the social sphere. Notably, two-week-old zebrafish stay on average 600-700 ms per entry in the social sphere, which is longer than the average duration of a swim bout (Fig. 2B).

### Two-week-old zebrafish proximity behavior is characterized by local adjustments of locomotion features

Our previous analyses showed that sociality greatly impacts overall locomotion (Fig. 2) and that specific features, such as turning behavior, vary significantly depending on inter-individual distance (Fig. 3C). We therefore investigated whether other features described in Fig. 2 vary along the d_HH_ axis and whether we could describe a set of characteristic locomotion properties occurring in the social sphere and at closer separations (i.e., d_HH_ < 15 mm), a set we will refer to as ‘proximity behaviors’ (Fig. 4). For clarity, we plotted data in the 0-22.5 mm range (full distributions including of control time-shifted data are available in Supp. Fig. 2, statistical relevant is reported in Supp. Table 1). We also highlighted average values at three relevant inter-fish distances (Fig. 4A; Table 2): one at very close proximity (d_HH_ = 3.5 mm), one within the social sphere (d_HH_ = 7.5 mm), and one outside the attention zone (d_HH_ = 35 mm). All locomotion features, with the exception of R bends (Suppl. Fig. 2A; Table 2, grey box), varied with head-head distance, while control time-shifted data were featureless.

**Figure 4.**
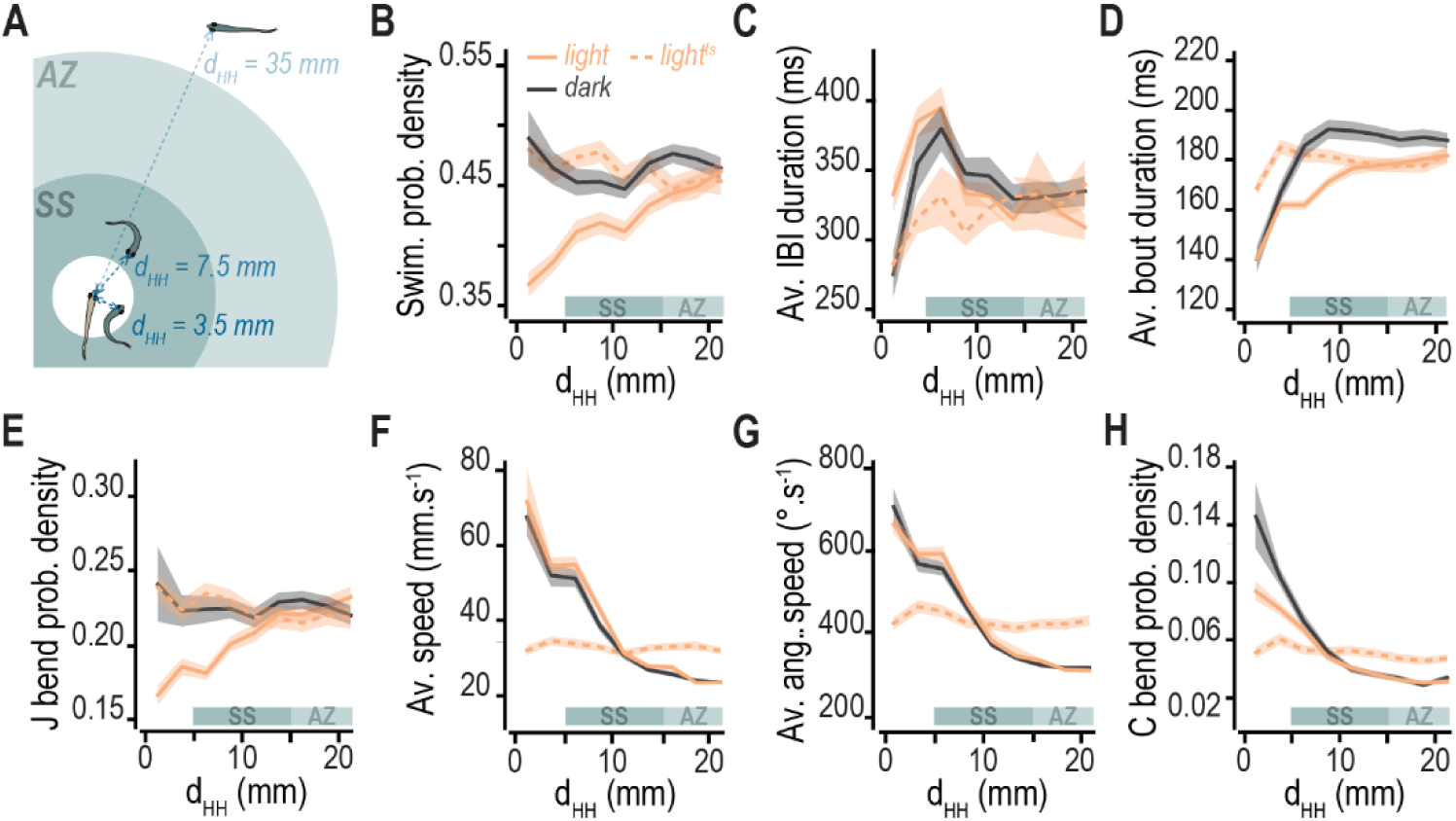
Proximity behavior is characterized by local adjustments of locomotion. **A**, We plot locomotion features across the range of head-head distance (d_HH_) and compare values of close proximity (d_HH_ = 3.5 mm), inside the Social Sphere (SS, d_HH_ = 7.5mm) or outside the Attention Zone (AZ, d_HH_ = 35mm, see median values and standard deviations in **Table 2**). Overall, locomotion features were similar across groups outside the Attention Zone (d_HH_ = 35 mm, see Supp. Fig. 2). Conversely, in the Social Sphere, some locomotion features were strongly modulated based on visual cues with decreased probability of swimming (**B**) correlated with increased average inter-bout interval (IBI, **C**) in parallel with decreased average bout duration (**D**), and decreased probability of J bends (**E**). Other features were strongly modulated at close proximity and independent of visual cues with increased average moving speed (**F**), increased average angular speed (**G**) and increased probability of C bends (**H**). Each individual dot represents a pair of fish. Traces represent the average across all pairs per condition, and the shaded area indicates the s.e.m.

**Table 2.** Social context impacts two-week-old zebrafish locomotion. Table presenting median values with standard variation for each locomotion feature observed in Fig. 4 and for R bends (Supp. Fig. 2). Data are presented for the three d_HH_ of interest: 3.5 (close proximity), 7.5 (in the Social Sphere) and 35 mm (outside the Attention Zone), and locomotion features are subcategorized depending on how they are affected along d_HH_: (i) in the Social Sphere, based on visual cues (orange box), (ii) in close proximity, independent of visual cues (blue box), and (iii) no effect (R bend, grey box). Statistical significance is reported in Supp. Table 1.

|  |  | Swimming | Inter-bout interval<br>*ms* | Bout duration<br>*ms* | J bend | Speed<br>*mm.s<sup>-1</sup>* | Angular speed<br>*°.s<sup>-1</sup>* | C bend | R bend |
| --- | --- | --- | --- | --- | --- | --- | --- | --- | --- |
| dark | 3.5 mm | 0.46 ± 0.11 | 337 ± 134 | 165 ± 28 | 0.20 ± 0.08 | 47.2 ± 18.5 | 581 ± 125 | 0.11 ± 0.05 | 0.17 ± 0.07 |
|  | 7.5 mm | 0.44 ± 0.07 | 335 ± 68 | 194 ± 27 | 0.22 ± 0.05 | 18.5 ± 9.24 | 466 ± 83.6 | 0.05 ± 0.03 | 0.16 ± 0.06 |
|  | 35 mm | 0.51 ± 0.07 | 303 ± 55 | 192 ± 28 | 0.23 ± 0.04 | 23.3 ± 2.59 | 306 ± 26.3 | 0.02 ± 0.01 | 0.16 ± 0.04 |
| light | 3.5 mm | 0.38 ± 0.06 | 375 ± 70 | 163 ± 13 | 0.20 ± 0.04 | 49.9 ± 13.6 | 589 ± 101 | 0.09 ± 0.03 | 0.16 ± 0.03 |
|  | 7.5 mm | 0.42 ± 0.07 | 335 ± 56 | 173 ± 14 | 0.21 ± 0.04 | 43.5 ± 9.72 | 480 ± 65.1 | 0.04 ± 0.02 | 0.16 ± 0.03 |
|  | 35 mm | 0.53 ± 0.07 | 269 ± 59 | 198 ± 23 | 0.26 ± 0.05 | 23.6 ± 1.94 | 324 ± 32.2 | 0.03 ± 0.02 | 0.17 ± 0.04 |
| dark<sup>ts</sup> | 3.5 mm | 0.51 ± 0.07 | 312 ± 72 | 194 ± 27 | 0.24 ± 0.06 | 27.7 ± 8.33 | 377 ± 83.5 | 0.04 ± 0.02 | 0.17 ± 0.05 |
|  | 7.5 mm | 0.51 ± 0.08 | 310 ± 68 | 194 ± 27 | 0.22 ± 0.07 | 27.0 ± 5.48 | 355 ± 61.9 | 0.04 ± 0.02 | 0.16 ± 0.04 |
|  | 35 mm | 0.47 ± 0.07 | 324 ± 70 | 189 ± 28 | 0.23 ± 0.06 | 26.4 ± 4.32 | 350 ± 47.2 | 0.04 ± 0.02 | 0.16 ± 0.04 |
| light<sup>ts</sup> | 3.5 mm | 0.48 ± 0.08 | 296 ± 60 | 183 ± 18 | 0.22 ± 0.06 | 34.8 ± 6.76 | 446 ± 102 | 0.05 ± 0.03 | 0.18 ± 0.04 |
|  | 7.5 mm | 0.47 ± 0.07 | 306 ± 52 | 179 ± 17 | 0.24 ± 0.05 | 31.8 ± 7.08 | 415 ± 83.9 | 0.05 ± 0.03 | 0.17 ± 0.03 |
|  | 35 mm | 0.49 ± 0.09 | 305 ± 57 | 178 ± 20 | 0.22 ± 0.05 | 32.5 ± 6.84 | 414 ± 74.9 | 0.05 ± 0.02 | 0.17 ± 0.03 |

Outside the attention zone, locomotion features were essentially similar across groups and also to control time-shifted data, indicating no effect of the presence of another individual at this large distance (Supp. Fig. 2; Table 2; Supp. Table 1). Conversely, we observed strong modulation of locomotion at closer range and could differentiate different patterns depending on proximity level. In the social sphere (d_HH_ = 7.5 mm), we found robust, visually driven modulation of zebrafish level and type of activity with fewer, shorter swim bouts and fewer J bends (Fig. 4B-E; Table 2, orange box). At closer range (d_HH_ = 3.5 mm), other features were more strongly impacted independently of visual cues with faster swim bouts, faster turning behavior and greater probability of C bends (Fig. 4F-H; Table 2, blue box), corroborating previous evidence of non-visually driven avoidance behavior at this distance range (Fig. 3C).

Overall, we found that proximity behavior is characterized by local modulation of locomotion and a general shift of strategy toward less frequent, shorter, faster bouts of swimming when zebrafish enter the social sphere. We show that this modulation is visually driven in the attention zone and the social sphere. In contrast, specific behaviors occurring in very close proximity, reminiscent of escape behavior, seemingly rely on other sensory systems, potentially the lateral line (Coombs, 2014; Groneberg et al., 2020).

### Two-week-old zebrafish engage in transient complex physical contacts

As discussed above, our analyses revealed strong signs of avoidance behavior occurring when fish are in close proximity (Fig. 3C,4F-H). We hypothesized that these avoidance responses could be triggered by social interactions involving physical contact. To test this hypothesis, we considered another measure of inter-fish distance, the closest distance (d_C_, Fig. 1C), allowing us to capture social interactions at a finer spatial scale.

Applying a similar analytical framework as for d_HH_, we first examined the average d_C_ per pair across the entire recording, then the probability density distribution for pairs in the light or dark, and included corresponding control time-shifted data (Fig. 5A-B). This analysis revealed similar broad patterns between lighting conditions as for d_HH_, with fish under illumination swimming on average closer than fish in the dark (Fig. 5A; median d_C_ = 16 mm ± 4 mm in the light versus 21 mm ± 4 mm in the dark, and 20 mm ± 4 mm in control time-shifted light data) and with a strong bias toward short distances (< 5 mm, Fig. 5B). Conversely, pairs in the dark do not show preference for any specific inter-fish distance (Fig. 5B). Note that both control time-shifted distributions display a high peak at d_C_ = 0, representing likelihood of fish intersecting, which almost never happens in real datasets due to the shallowness of the arenas (see Methods).

**Figure 5.**
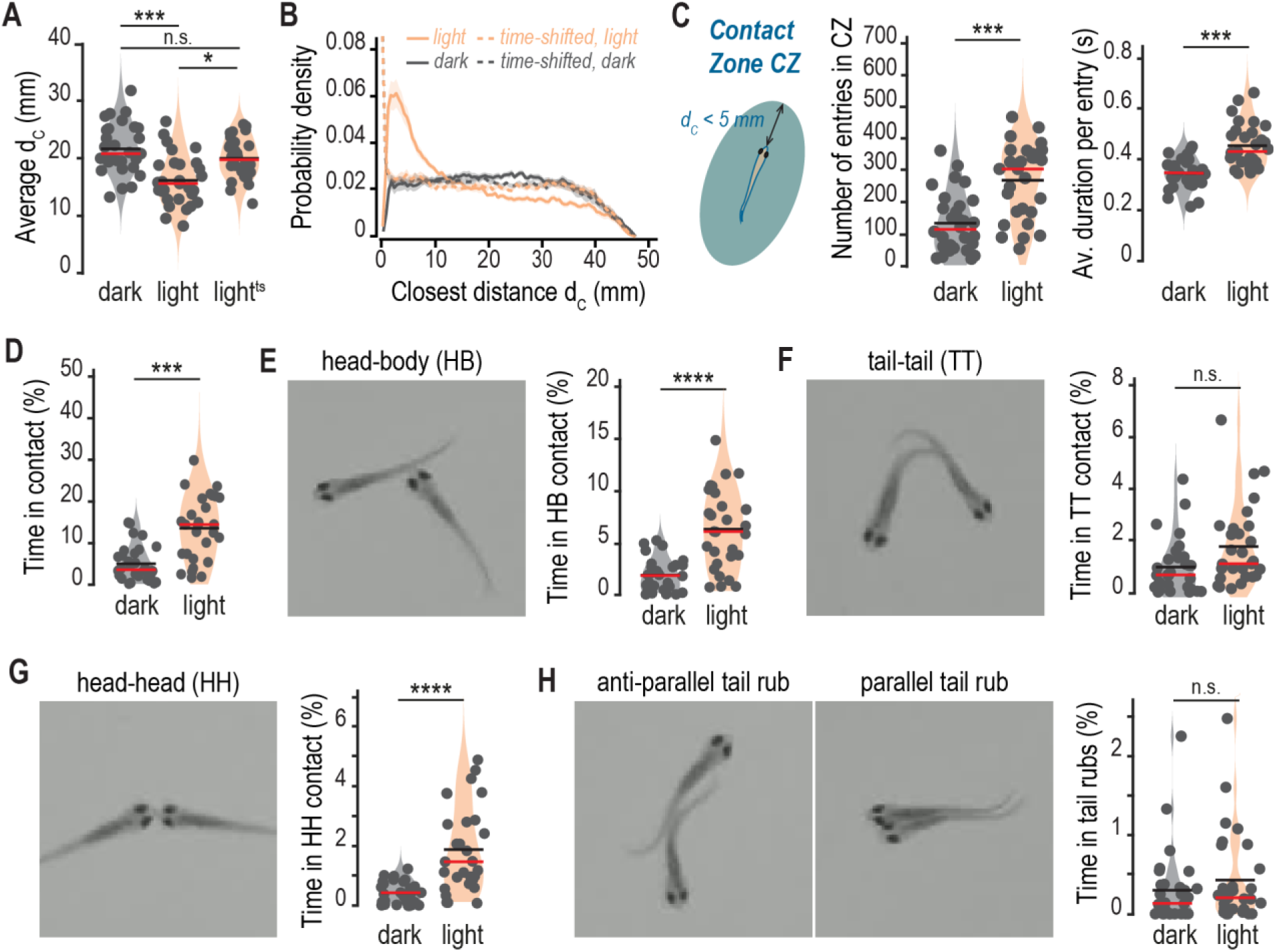
Two-week-old zebrafish engage in diverse transient physical contacts. The average closest distance (d_C_) between two fish is significantly reduced in lit arenas compared to arenas in the dark (**A**) and the overall probability distribution is skewed toward short distances (**B**). **C**, The Contact Zone (CZ) is the zone of high probability for d_C_ (< 5 mm, left schematic). Pairs in the light enter (middle graph) and stay (right graph) in the contact zone significantly more than pairs in the dark. Average fraction of moving time that zebrafish spend in contact (**D**), further subdivided into head-body contacts (HB, **E**) and tail-tail contacts (TT, **F**), and then head-head contacts (HH, **G**) and parallel or anti-parallel tail rubs (**H**). Each individual dot represents a pair of fish. Red and black bars in graphs indicate the median and the mean, respectively. Statistical significance was tested using two-sample Kolmogorov-Smirnov tests.

We postulated that animals are in a position to engage in physical contact when the closest distance is shorter than 5 mm. We therefore defined the ‘contact zone’ (CZ) for d_C_ < 5 mm (also true if d_HH_ < 5 mm) (Fig. 5C, left schematic) and quantified how often and for how long zebrafish enter this zone. Under illumination, zebrafish enter the contact zone significantly more often (Fig. 5C, middle panel; median number of entries = 304 ± 116 in the light versus 114 ± 93 in the dark) and for longer periods of time (Fig. 5C, right panel; median duration = 432 ms ± 87 ms in the light versus 346 ms ± 60 ms in the dark) than fish in the dark. It should be noted that these durations are comparatively shorter than the ones observed for the social sphere (Fig. 3G), highlighting different dynamic features between the two zones.

We next sought to determine the types of social interactions occurring in the contact zone and whether they involve engagement in physical contact. We first manually analyzed portions of recordings where our analysis pipeline indicated that fish were in the contact zone. This analysis revealed that young zebrafish engage in an unexpectedly diverse range of physical contacts. We established a human-curated list of these contacts based on our manual observations and implemented their automatic detection using a combination of objective geometric criteria, such as inter-fish distance, relative orientation, and number and position of points of contact on the body (see Methods). Incorporating this detection into our automated analysis pipeline allowed the classification of physical interactions into broad, biologically meaningful categories and unbiased quantification of the number, duration and frequency of events for each type of contact motif.

Our analysis revealed that, overall, young zebrafish spend a substantial amount of time engaged in physical contact, independent of the type of motif, and that this behavior is largely dependent on visual cues (Fig. 5D; median percentage of moving time spent in contact = 14.4% ± 8% in the light versus 3.6% ± 4% in the dark). We distinguished between two main categories of physical contacts: head-body contacts (Fig. 5E; Suppl. Movie 1) and body-body or tail-tail contacts (Fig. 5F; Suppl. Movie 2). Head-body contacts constituted the main type of contact displayed in our arenas, occurring three times as often as tail-tail contacts (median percentage of time spent in head-body contacts = 6.1% ± 3.8% in the light versus 1.9% ± 1.5% in the dark; median percentage of time spent in tail-tail contacts = 1.1% ± 1.6% in the light versus 0.7% ± 1.1% in the dark). Within head-body contacts, we further isolated specific events when the two fish enter in contact through both their heads, head-head contacts, which only represented a small fraction of total head-body contacts (Fig. 5G; Suppl. Movie 3; median percentage of time spent in head-head contacts = 1.5% ± 1.4% in the light versus 0.4% ± 0.4% in the dark). In the same vein, we described tail rubs within the tail-tail contact category. Although rare, these events of parallel or anti-parallel tail-tail contacts, were remarkable for the high degree of coordination between two individuals they seemingly require (Fig. 5G; Suppl. Movies 4 and 5; median percentage of time spent in tail rubs = 0.2% ± 0.6% in the light versus 0.1% ± 0.5% in the dark). Importantly, these distinctions enabled us to separate unilateral, asymmetrical interactions, initiated by one fish and imposed on the other (for example, a head-body contact), from mutually coordinated, reciprocal ones, that assume some degree of reciprocity in the behavior (for example, head-head contacts or tail rubs).

Combined, our observations drawn from the analysis of closest distance distributions and the definition of the contact zone established that two-week-old zebrafish frequently engage in a diversity of transient physical contacts that are reminiscent of complex behaviors typically described in the adult social repertoire and that can be either symmetrical or asymmetrical in nature (Kalueff et al., 2013, 2025). Notably, all described contact motifs were drastically reduced in the dark, in the absence of visual cues, emphasizing the importance of the visual system for two-week-old zebrafish to engage in these physical interactions. Most physical interactions in our dataset also appeared to be non-reciprocal, with a majority of head-body contacts compared to other types of contacts, and are therefore more likely to trigger startle or escape responses in the subjected fish. This result was consistent with our previous observations that zebrafish exhibit strong avoidance behavior in the contact zone range, and with our hypothesis that this avoidance behavior is mediated by physical encounters.

## Discussion

In this study, we set out to characterize the behavior of freely interacting two-week-old zebrafish, examining a stage when social behavior is just emerging (Stednitz & Washbourne, 2020). By developing a hybrid analysis pipeline that automatically extracts human-identified motifs using behavioral data from entire recordings, independent of swimming structure, we were able to uncover previously undescribed features of early zebrafish social behavior. Our approach proved particularly valuable in disentangling genuine social dynamics from artifacts imposed by arena geometry and thigmotaxis behavior, and in uncovering different tiers of locomotion modulation depending on inter-fish distance. As a result, we obtained an integrated and interpretable picture of the emergence of social behavior, which had not been revealed previously through either purely supervised or purely unsupervised methods (Stednitz et al., 2025).

The importance of the visual system for zebrafish behavior, especially social behavior, has been well documented in the literature (Dreosti et al., 2015; Engeszer et al., 2004, 2007; Stednitz et al., 2025). We therefore compared two conditions throughout our analyses, with zebrafish freely swimming either under homogenous illumination or in the dark, allowing the delineation of motifs that are mediated by visual cues from those that are not. In single individuals, we described a robust, light-induced increase in locomotion activity levels and speed while other features, such as turning or thigmotaxis behaviors, remained constant in different lighting conditions. This suggests that pathways that modulate bout duration, frequency and speed may integrate different sensory cues, including visual cues, than the ones determining bout type or general spatial preference. Interestingly, light-induced activation of locomotion level and speed was dampened in a social context, suggesting that zebrafish may change locomotion strategy to adjust exploration when in presence of another individual. Notably, the influence of visual cues on locomotion becomes more nuanced and depends on inter-fish distance, integrating information about both lighting condition and visual stimuli represented by the other individual.

We propose a conceptual framework to summarize the collective behavioral features displayed at various inter-fish distances, defining four relevant social spaces, as illustrated in the Graphical abstract: 1/ outside the attention zone, zebrafish are less or unresponsive to the presence of another individual; 2/ the attention zone, where, based on long-range visual cues, zebrafish start showing reduced stochastic movements and biased turning behavior toward their counterpart; 3/ the social sphere, resulting from the social biasing of turning behavior, where zebrafish linger and adopt less frequent, shorter but faster bouts of swimming; 4/ the contact zone, where zebrafish transiently engage in physical interactions that, in turn, trigger robust avoidance behavior. While the initiation of physical contacts relies strongly on visual cues, avoidance depends on other, non-visual, short-range sensory cues, most likely mediated by the lateral line system as in stereotypical touch-escape response (Kohashi & Oda, 2008).

One of our central findings is that two-week-old zebrafish actively maintain a preferred range of inter-individual distances through visually mediated biased turning behavior, creating a space that we named the ‘social sphere’. Identifying interaction rules for animals and explaining how collective behaviors such as shoaling, flocking, and swarming can arise is a major topic of current investigation (Ballerini et al., 2008; Couzin & Krause, 2003; Hinz & Polavieja, 2017; Sumpter, 2006). As noted earlier, zebrafish are a useful model for such studies, especially at stages when social interactions are developing. The attraction of pairs of roughly two-week-old zebrafish to each other has been characterized previously (Dreosti et al., 2015; Engeszer et al., 2004, 2007; Stednitz et al., 2018, 2025; Stednitz & Washbourne, 2020), notably in pioneering work by Hinz and de Polavieja (Hinz & Polavieja, 2017), who quantified the reduction in pair separation and preferential turning toward neighbors as early as 12 dpf. From these observations, the authors built a model in which attraction is probabilistic in time. With the attraction probability as a parameter, the model could mirror the observed changes in mean inter-fish distance with fish age, though the full probability distribution of inter-fish distances, a stringent test as we note here, was not modeled. More intriguingly, this and other work that has reported inter-fish attraction have left open the question of *how* attraction occurs.

The model we introduce here reproduces the shape of the full distribution of inter-fish distances and by examining its components, which include socially induced changes in bout duration and speed, we find that a surprisingly dominant driver is the socially mediated variance of turning angle. Namely, over a range of inter-fish distances spanning about two to four average body lengths and orientations corresponding to the other fish being forward, the standard deviation of the turning angles the fish adopts is considerably reduced, focusing the trajectory into more persistent paths. This mechanism is somewhat reminiscent of klinokinesis, i.e., a change in the frequency of turning in response to stimuli (Fraenkel & Gunn, 1961). Klinokinesis is known to occur in contexts as diverse as bacterial chemotaxis via modulation of ‘tumbles’ (Howard, 1993) and aggregation of the nematode *Caenorhabditis elegans* via density-dependent turning rates (Ding et al., 2019). Whether it occurs in fish schools or insect swarms, separate from directional steering toward neighbors, remains unclear. Regardless, socially modulated turn variance as we observe in our data is distinct from klinokinesis, as the breadth of possible turns can be tuned independently from the frequency of turning, and we suggest instead that it may be a general mode of animal behavioral control that can be investigated in other systems. How the dyadic behavioral observations and model we describe here develop into more complex forms with increased animal number and age will be important to investigate, possibly shedding light on the ontogeny of complex social and collective behavior.

We also describe how, once in the social sphere, zebrafish adopt a different mode of locomotion characterized by less frequent, shorter but faster bouts of swim and increased likelihood of J bending. The case of J bends was interesting among locomotion features dynamically adjusted depending on inter-individual distance. J bending is typically associated with J turns, a behavioral motif originally described as part of the stereotyped prey capture sequence in zebrafish larvae (Budick & O’Malley, 2000b; Marques et al., 2018; McElligott & O’Malley, 2005). In prey capture, J turns enable zebrafish to re-orient themselves to align with their prey and precede darting forward to catch this prey. Our observations therefore suggest a broader role of J bending that could be deployed in contexts other than prey capture that also require fine directional adjustment toward salient targets, for example here as part of approaching behavior during social interactions. This observation is in line with growing appreciation that discrete locomotor motifs in larval zebrafish repertoire can serve multiple behavioral functions depending on context (Johnson et al., 2020; Marques et al., 2018) and raises the question of how similar motor programs can be recruited through different sensory stimuli.

Finally, we describe for the first time a set of complex, physical interactions among two-week-old zebrafish, taking the form of transient, quick contacts accompanied by robust avoidance responses. Notably, our analysis reveals that two-week-old zebrafish already exhibit a diversity of contacts that are not all equivalent: while some patterns appear cooperative, mutually initiated, and require a substantial degree of coordination (e.g., tail rubs), others are asymmetrical or non-cooperative (e.g., head-body contacts). This heterogeneity echoes the diversity of contact patterns described in adult zebrafish, including lateral approaches, nudges, and behavioral sequences of both positive and negative valence associated with affiliative and agonistic interactions (Kalueff et al., 2013, 2025; Pham et al., 2012). This observation suggests that zebrafish apply partner-contingent behavioral adjustments from the onset of sociality and that complex social repertoires are established earlier than previously appreciated. It also raises important questions about the biological function of physical contacts at this age. Physical contacts between conspecifics could facilitate the transfer of chemical or microbial cues, consistent with evidence that skin-associated microbiota and olfactory signals play roles in social recognition and group cohesion in fish (Reverter et al., 2018). These patterns might also serve a social learning function, allowing naive animals to acquire and refine social skills and associated motor patterns that they will use as adults and will be critical for survival (Huzard et al., 2022; Yu et al., 2022). Finally, physical contacts might contribute to social buffering and enhance overall well-being, as has been documented in mammals and adult zebrafish (Faustino et al., 2017; Liu & Yuan, 2016). Distinguishing among these possibilities will require targeted experimental manipulations.

Together, our results establish foundational description of two-week-old zebrafish social behavior and demonstrate that they are active social agents displaying behavioral patterns that are precursors of adult social interactions. The analytical framework we introduce offers a straightforward, interpretable picture of sociality while being sensitive enough to detect dynamic features of social interactions and to distinguish between interactions of different valence. Critically, our study opens a new window into subtle shifts in social behavior that may accompany the early stages of neuropsychiatric or neurodevelopmental conditions in zebrafish disease models.

## Resource availability

All analysis code and simulation software is available in the public GitHub repository https://github.com/RaghuParthasarathy/Zebrafish_social_behavior_analysis_June2026.

## Acknowledgments

The authors thank the University of Oregon Aquatic Animal Care Services staff for assistance maintaining fish lines and rearing larvae to two weeks of age. We also thank Luca Mazzucato, Eric Corwin, Greg Stephens, and members of the Guillemin, Eisen and Parthasarathy laboratories for insightful discussions and invaluable feedback on the project and the manuscript. Funding for this project was acquired by KG, JE and RP (NIH 1RM1GM158513-01), and by JE (Gordon and Betty Moore Foundation, Symbiosis in Aquatic Systems Initiative, award 9205). This manuscript is the result of funding in part from the National Institutes of Health (NIH). It is subject to the NIH Public Access Policy. Through acceptance of this federal funding, NIH has been given a right to make this manuscript publicly available in PubMed Central upon the official date of publication, as defined by NIH.

## Author contributions

LD, KG, JE and RP conceptualized the study. Zebrafish behavior experiments were conducted by LD. Analysis software was written by RP. Manual checking of behavioral annotations was undertaken by LD. The behavior model and its software implementation were developed by RP. Analysis and interpretations were made jointly by LD and RP. Figures and supplementary elements, including movies, were prepared by LD. The manuscript was written by LD and RP, reviewed by JE and KG.

## Declaration of generative AI and AI-assisted technologies

Artificial intelligence assisted with generating some of the analysis code, especially related to data import and export, slicing datasets based on various conditions and constraints, and plotting data. Artificial intelligence was also used to interactively develop simulation code, especially related to plotting, testing, interrogation of modeling choices, and iteration of various models. AI assistance was provided by Claude, specifically the 4.x Sonnet and Opus models.

## Supplemental information titles and legends

See attached document.

## Methods

### Ethics statement

All zebrafish experiments were approved by the University of Oregon Institutional Animal Care and Use Committee (protocol 20-15).

### Zebrafish lines and husbandry

All zebrafish lines were maintained according to the Zebrafish Book guidelines at 28°C under a 14/10 light/dark cycle. Behavioral experiments were conducted using ABC wild-type fish raised to two weeks.

### Behavior recording

Clutches were collected from pairs of ABC adult fish and kept separate. At day 0, 200 eggs were selected from each clutch, bleached, then subdivided into four dishes of 50 in fresh embryo medium. At 4 days post fertilization (dpf), the 25 best looking larvae with inflated swim bladders were selected from each dish and transferred to the nursery in separate tanks where they were fed and monitored daily until experimental sampling. At 14 dpf, animals were retrieved from the nursery and checked for growth. Only animals of standard length 7 mm and above were tested for social behavior (Stednitz & Washbourne, 2020). We adapted a dyad assay from previous publications (Bruckner et al., 2022; Stednitz et al., 2018) to record the behavior of freely swimming and interacting animals. Briefly, animals were placed in shallow circular chambers (diameter 5 cm, depth 2 mm) carved in an acrylic plate placed atop an infra-red panel (EnvironmentalLights) and filled with fresh system water. After 5 minutes of habituation, animals were recorded for 10 minutes from the top at 25 Hz using a monochromatic camera (BFS-U3-88S6M-C USB 3.1 Blackfly S, Edmund Optics #11-511) equipped with a 35-mm focal lens (Edmund Optics #63-247) and an infra-red bandpass filter (Edmund Optics #84-802). After experimental testing, all fish were humanely euthanized in ice-cold water according to standard protocols. Chambers were thoroughly rinsed with system water between each trial. All behavioral experiments were performed in a room kept at 28°C and 50% humidity with the behavioral setup placed on an air table to avoid disruptive vibrations.

### Behavior analysis

Zebrafish body positions in all movies were assessed using ZebraZoom software (https://zebrazoom.org/ (Mirat et al., 2013)), which processes each frame and outputs the x and y coordinates of 10 points along the body of each fish. The resulting arrays of coordinates were processed using custom Python code to extract parameters describing basic features of locomotion such as speed, tail bending, and bout duration, as well as complex features of social behavior such as inter-fish distance and relative orientation.

We first identified and repaired tracking errors: 1/ we found the anterior-most head position of each fish to be unreliable and recalculated it with a linear extrapolation based on body positions 2 through 4; 2/ we identified frames in which ZebraZoom failed to identify the proper number of fish, recording these as ‘bad’ frames to exclude from further analysis; 3/ in an attempt to improve tracking of fish identity, we also re-calculated the linkage of fish across frames, i.e., the identification of which fish in a given frame corresponds to the same fish in the next frame, using all the body positions and minimizing the L2 norm of the Euclidean distance between body positions across adjacent frames.

Next, we evaluated single fish position and locomotion properties at each frame. These consist of the distance from the center of the circular arena; the speed, calculated from the frame-to-frame displacement of the head position; the turning angle and angular speed, calculated from the frame-to-frame change in heading angle, itself calculated based on the frame-to-frame change in direction vector based on the two most anterior body positions; the tail bending angle, calculated as the angle between the best-fit lines to the anterior and posterior halves of the fish; and the occurrences of J-, R-, and C-bends, defined as in prior literature for tail bending angles 10-50°, 50-100°, and > 100°, respectively. Additionally, for speed and angular speed, an average value was calculated per pair and recording by averaging single values from all frames where fish were detected as ‘moving’, defined here as when fish displacement was over a given threshold (9 mm/s).

For datasets with two fish, our program then assessed basic features of the pair’s configuration in each frame. These include the inter-fish distance, both the distance between head positions and the closest distance between any points, and the relative orientation of each fish with respect to the other (Fig. 1F). Note that the relative orientation is the angle between the heading vector of a fish and the vector connecting its head and that of the other fish (as in (O’Shaughnessy et al., 2024)); |ф_i_| < 90° corresponds to fish *i* facing towards the other fish, and ф_i_ < 0 corresponds to the other fish being to the left of fish *i*.

We identified specific behavioral motifs for pairs of zebrafish based on temporal and geometric properties. As above, denoting the heading angle of fish *j* as Ɵ_j_, the relative orientation to the other fish as ф_i_, and the closest distance between the fish as d_C_, these behavioral motifs include: ‘contact’, when d_C_ < 1.3 mm independent of duration; ‘tail rubbing’, when the two fish are oriented parallel or antiparallel with at least two of the four most posterior tail positions within 2.0 mm of each other for at least two frames (80 ms), and cos(Θ_1_ – Θ_2_) < −0.8 or > 0.8 for antiparallel or parallel tail rubs, respectively; and ‘staying in the social sphere’ and ‘staying in the contact zone’ when the two fish maintain 5 < d_HH_ < 15 mm or 0 < d_C_ < 5 mm, respectively. For all these behavioral motifs, the frames in which the behaviors are detected are recorded along with the number of events and their durations. We calculated the fraction of time spent in these motifs by dividing the recorded number of frames for a given motif by the total number of frames in which any of the two fish is moving (according to the threshold speed of 9 mm/s), allowing us to normalize per level of activity.

### Simulation of zebrafish behavior

Our simulation of zebrafish motion is described in detail in the Supp. Text 1. In brief, the trajectory of each fish is modeled as a biased random walk, with steps corresponding to bouts and pauses corresponding to inter-bout intervals. Kinematic parameters such as bout length and duration are jointly sampled from the data, binned by radial position (r) and inter-fish distance (d_HH_). The turning angle at each step is determined by 1/ calculating a weighted circular mean Δ*θ*_M_ of the mean turning angle from single-fish data at the current (r, ψ) bin, where ψ is the wall alignment angle, and a social turning angle toward the neighbor given by the relative orientation angle (ф), with the distance-dependent weight calculated from the observed pair turning data; 2/ sampling a random turning angle Δ*θ*_0_ from single-fish data, scaling its difference from the mean single-fish turning angle by *f*, the ratio of the pair turns’ standard deviation in each (d_HH_, |ф|) bin to its long-distance asymptote, and adding the result to Δ*θ*_M_. In other words, we shift the single-fish turn distribution by a distance-dependent social weight and rescale its width by a distance- and orientation-dependent focusing of variance. The simulation is then run, with reflective boundary conditions at the circular arena walls, until the simulated time reaches the experiment duration, and properties such as inter-fish distance distributions are calculated.

