## Supplementary Figures and Text for "Social modulation of activity and orientation fosters complex interactions in young zebrafish"

**Supplementary Figure 1**

**Supplementary Figure 2**

**Supplementary Table 1**

**Supplementary Text 1**

We describe in detail the model of zebrafish pair behavior sketched in the main text, including comparisons to experimental data and assessment of the importance of various model components. This text contains Supplementary Figures 3 through 15.

**Supplementary Movie 1**

Illustrative movie of a head-body contact as illustrated in Fig. 5E. The movie spans 1.2 sec and is slowed down 5x. Scale bar, 3 mm.

**Supplementary Movie 2**

Illustrative movie of a tail-tail contact as illustrated in Fig. 5F. The movie spans 1.24 sec and is slowed down 5x. Scale bar, 3 mm.

**Supplementary Movie 3**

Illustrative movie of a head-head contact as illustrated in Fig. 5G. The movie spans 0.72 sec and is slowed down 5x. Scale bar, 3 mm.

**Supplementary Movie 4**

Illustrative movie of an anti-parallel tail rub as illustrated in Fig. 5H, left picture. The movie spans 1.52 sec and is slowed down 5x. Scale bar, 3 mm.

**Supplementary Movie 5**

Illustrative movie of a parallel tail rub as illustrated in Fig. 5H, right picture. The movie spans 3.0 sec and is slowed down 5x. Scale bar, 3 mm.

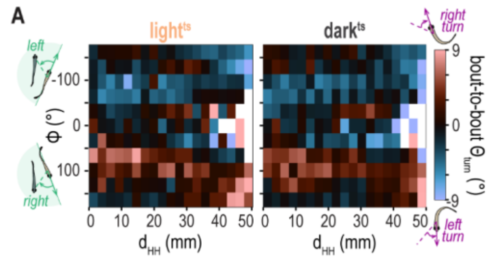


**Supplementary Figure 1.**
Two-dimensional plots of the average bout-to-bout turning angle ($\Theta_{turn}$) as a function of head-head distance ($d_{HH}$) and relative orientation ($\phi$) for control, time-shifted dataset ligh$^{ts}$ and dark$^{ts}$, matching the presentation of the same maps for light and dark datasets in Fig. 3C. White squares indicate bins with no data.

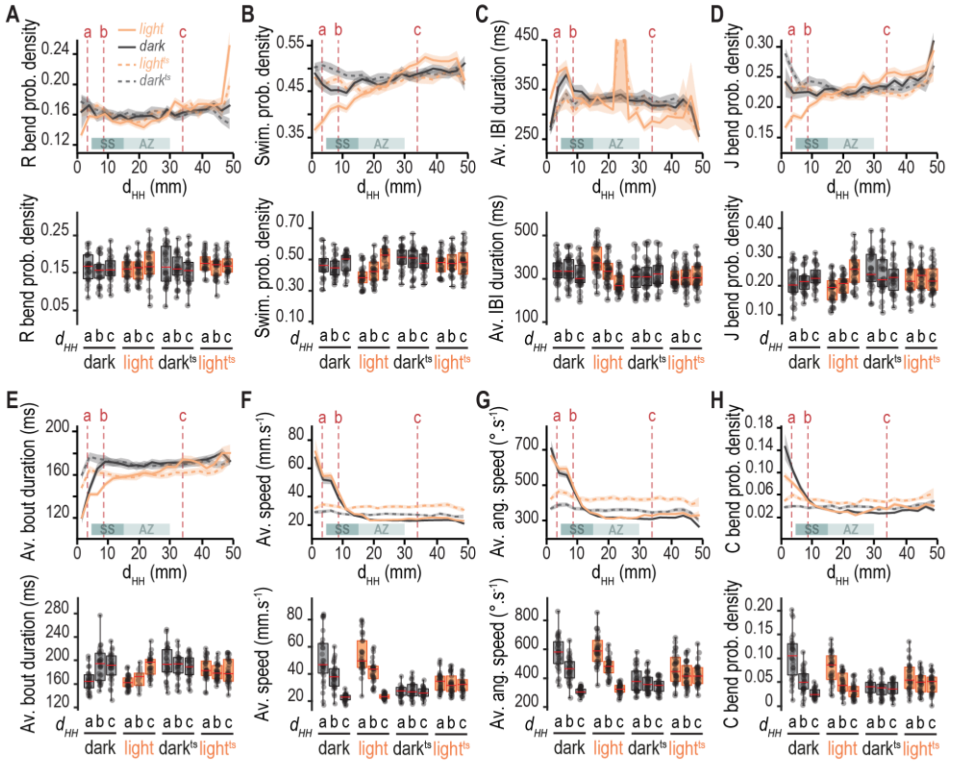


**—Supplementary Figure 2.**

Full distributions of locomotion features across the range of head-head distance ($d_{HH}$) as presented in Fig. 4 with the addition of R bends (**A**). We measured and compared values in close proximity (a, $d_{HH}$ = 3.5 mm), inside the Social Sphere SS (b, $d_{HH}$ = 7.5 mm) or outside the Attention Zone AZ (c, $d_{HH}$ = 35 mm). Individual values are presented with box plots under corresponding distributions. Median and standard deviation values across individuals from each group at each $d_{HH}$ of interest are presented in Table 2. Each individual dot represents a pair. Traces represent the average across all pairs per condition, and the shaded area indicates the s.e.m. Statistical significance was tested using two-sample t-tests without correction for multiple comparisons and is presented in Supplementary Table 1.

| | | Dark | | Dark$^{ts}$ | | Light | | Light$^{ts}$ | | | | |
|---|---|---|---|---|---|---|---|---|---|---|---|---|
| | | SS | Far | SS | Far | SS | Far | SS | Far | Dark vs Light | Dark vs Dark$^{ts}$ | Light vs Light$^{ts}$ |
| Prob. Density R bend | CZ | 0.54915 | 0.50315 | 0.50154 | 0.29360 | 0.71969 | 0.18990 | 0.98666 | 0.78900 | 0.29449 | 0.92782 | 0.42716 |
| | SS | X | 0.97498 | X | 0.62282 | X | 0.31586 | X | 0.75897 | 0.91166 | 0.95762 | 0.52414 |
| | Far | X | X | X | X | X | X | X | X | 0.45722 | 0.65110 | 0.82140 |
| Prob. Density Swimming | CZ | 6.31E-01 | 3.26E-01 | 4.85E-01 | 1.65E-01 | 6.68E-02 | 5.85E-09 | 4.55E-01 | 4.79E-01 | 0.00120 | 0.08467 | 0.00020 |
| | SS | X | 6.72E-02 | X | 5.20E-01 | X | 1.06E-05 | X | 9.66E-01 | 0.07365 | 0.05269 | 0.00218 |
| | Far | X | X | X | X | X | X | X | X | 0.24688 | 0.65362 | 0.14891 |
| Average IBI | CZ | 9.15E-01 | 2.25E-01 | 7.73E-01 | 3.99E-01 | 2.92E-03 | 2.75E-07 | 4.98E-01 | 8.34E-01 | 0.22753 | 0.20023 | 0.00015 |
| | SS | X | 6.89E-02 | X | 5.64E-01 | X | 2.88E-03 | X | 6.36E-01 | 0.38358 | 0.11725 | 0.05192 |
| | Far | X | X | X | X | X | X | X | X | 0.04128 | 0.46283 | 0.09629 |
| Prob. Density J bend | CZ | 9.27E-01 | 5.38E-01 | 4.34E-01 | 1.05E-01 | 1.64E-01 | 1.03E-06 | 6.29E-01 | 9.04E-01 | 0.03161 | 0.08092 | 0.00593 |
| | SS | X | 4.71E-01 | X | 4.78E-01 | X | 2.93E-05 | X | 5.09E-01 | 0.04495 | 0.25941 | 0.01530 |
| | Far | X | X | X | X | X | X | X | X | 0.07832 | 0.84632 | 0.01278 |
| Average Bout Duration | CZ | 2.02E-03 | 2.77E-03 | 6.83E-01 | 2.64E-01 | 1.21E-02 | 1.11E-08 | 4.10E-01 | 5.00E-01 | 0.22094 | 0.00022 | 0.00000 |
| | SS | X | 9.29E-01 | X | 4.97E-01 | X | 2.13E-05 | X | 9.42E-01 | 0.00051 | 0.79147 | 0.01389 |
| | Far | X | X | X | X | X | X | X | X | 0.67154 | 0.77963 | 0.03049 |
| Average Speed | CZ | 9.10E-04 | 1.35E-11 | 4.93E-01 | 1.97E-01 | 3.80E-04 | 0.00E+00 | 3.66E-01 | 4.41E-01 | 0.54473 | 0.00000 | 0.00000 |
| | SS | X | 3.01E-12 | X | 4.44E-01 | X | 1.00E-14 | X | 8.84E-01 | 0.08825 | 0.00000 | 0.00003 |
| | Far | X | X | X | NaN | X | X | X | X | 0.10421 | 0.00004 | 0.00000 |
| Average Angular Speed | CZ | 3.24E-04 | 1.00E-15 | 2.76E-01 | 1.98E-02 | 2.36E-06 | 0.00E+00 | 1.09E-01 | 6.89E-02 | 0.43110 | 0.00000 | 0.00001 |
| | SS | X | 2.17E-13 | X | 1.43E-01 | X | 1.00E-15 | X | 8.62E-01 | 0.48767 | 0.00001 | 0.01264 |
| | Far | X | X | X | X | X | X | X | X | 0.01776 | 0.00041 | 0.00000 |
| Prob. Density C bend | CZ | 1.35E-06 | 6.12E-12 | 4.82E-01 | 4.33E-01 | 7.79E-06 | 3.42E-08 | 2.31E-01 | 6.09E-02 | 0.03324 | 0.00000 | 0.01029 |
| | SS | X | 2.55E-05 | X | 9.97E-01 | X | 3.72E-02 | X | 5.09E-01 | 0.53250 | 0.01658 | 0.50232 |
| | Far | X | X | X | X | X | X | X | X | 0.15032 | 0.01372 | 0.03348 |

**Supplementary Table 1.**
Statistical significance between each group and $d_{HH}$ of interest presented in Supp. Fig. 2 was tested using two-sample t-tests without correction for multiple comparisons. P-values are provided in this table.

### Supplementary Text 1 - Behavior Model

#### Summary

Because larval zebrafish locomotion takes the form of bouts of motion punctuated by inter-bout intervals of rest, we model fish trajectories as biased random walks of linear constant-velocity steps. We sample parameters such as bout length and duration from experimental data and incorporate a social bias in turning angles directed toward the neighbor and a socially modulated reduction in the variance of turning angles. Notably, all the model parameters, including the turning bias weight and the variance reduction, are derived from the data; **there are no free (adjustable) parameters.** Simulations using the model quantitatively reproduce the inter-fish distance distribution $p(d_{HH})$ for pairs of fish in lit circular arenas and pairs in the dark; see Fig. 3F in the main text and Supp. Fig. 3 below, noting especially the apparent preference for separations around 5-15 mm. In addition, the simulated distributions of the fishes' radial position distributions also match the data (Supp. Fig. 4 below) As shown in the following text, alteration of the model assumptions or omission of its components clearly worsens the match between simulations and experimental data. The model is memoryless, considering only instantaneous decisions by the fish, and its success at describing observations suggests that memory or persistent behavioral states may be absent or relatively unimportant at this developmental stage.

Commented [JE1]: see previous comment

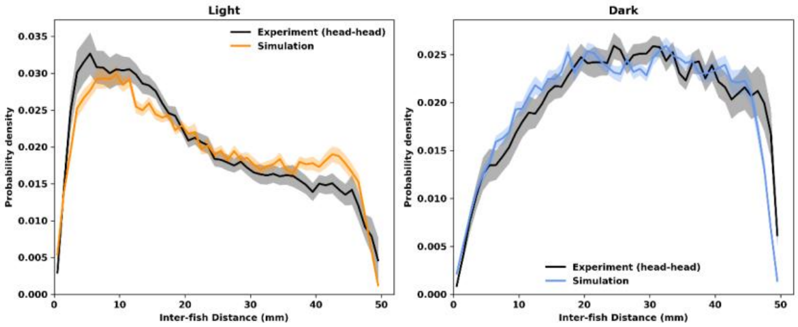


**Supplementary Figure 3.**
The probability distribution of inter-fish head-to-head distances, $p(d_{HH})$, for pairs of fish in circular arenas of radius 25 mm in (left panel) light and (right panel) dark. The experimental data, as discussed in the main text, are drawn from 39 and 38 datasets of 10-minute duration each in lit and unlit arenas, respectively. For each condition, we generated 40 simulated 10-minute trajectories. Solid lines show the mean across datasets or simulations, with the shaded bands indicating the s.e.m.

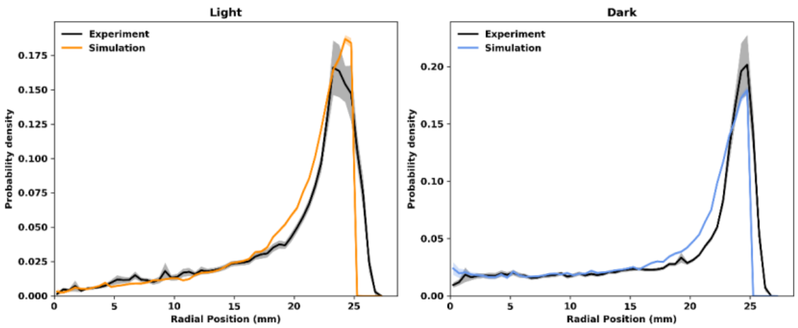


**Supplementary Figure 4.**
The probability distribution of radial position, p(*r*), for pairs of fish in circular arenas of radius 25 mm in (left panel) light and (right panel) dark, with experimental and simulation data plotted as in Supp. Fig. 3. The radial distribution function is area-normalized by 1/r.

### Sampling bout properties from single-fish experimental data

Larval zebrafish locomotion takes the form of bouts of activity punctuated by inter-bout-intervals of rest. From tracking data from experiments using single fish in circular arenas, we identify bouts and inter-bout intervals. For each inter-bout interval (IBI) *j*, we tabulate the mean coordinates (radial coordinate $r_j$ and polar angle $\gamma_j$), duration $T_j$, and direction of the displacement of the mean position into this IBI from the previous IBI ($\theta_j$). Considering the bout that follows IBI *j*, we tabulate the bout duration ($\Delta t_j$), displacement magnitude or step size ($\Delta s_j$), and turning angle $\Delta\theta = \theta_{j+1} - \theta_j$. These are binned by *r* and ψ, where ψ is the wall alignment angle, defined as $\psi = \theta - \gamma$. This ψ describes the heading direction relative to the wall normal; ψ = 0 indicates a step directed radially outward and ψ = ± 90° indicates a trajectory directed along the wall (Supp. Fig. 5).

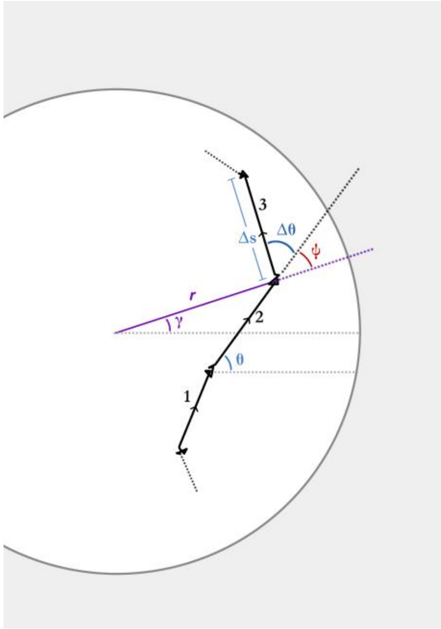


**Supplementary Figure 5.**
Schematic diagram illustrating a series of bouts (black lines, numbered) in a circular arena and the notation used in the analysis, specifically polar coordinates (r, γ), bout displacement (Δs), bout direction angle (θ) and change in angle (Δθ), and wall alignment angle (ψ).

**Commented [LD2]:** Maybe number them?

**Commented [RP3R2]:** Done.

### Simulating single fish motion

We simulate single-fish motion as a memory-less random walk with linear steps of magnitude (Δs), turning angle (Δθ), duration (Δt), and inter-bout interval duration (T) drawn from r- and ψ-binned experimental values. We apply reflective boundary conditions at the circular arena wall. Using the single-fish-derived parameters reproduces the observed single fish radial probability distribution, p(*r*), extremely well. This matching between experiment and simulation is not trivial; neglecting the wall alignment angle ψ or mis-characterizing the relationship between wall angle and the current heading angle, for example, leads to poor agreement. Simulating single fish motion by sampling from experiments with fish in light reproduces the p(*r*) from fish in light; sampling from experiments in the dark reproduces the p(*r*) from fish in the dark (Supp. Fig. 6).

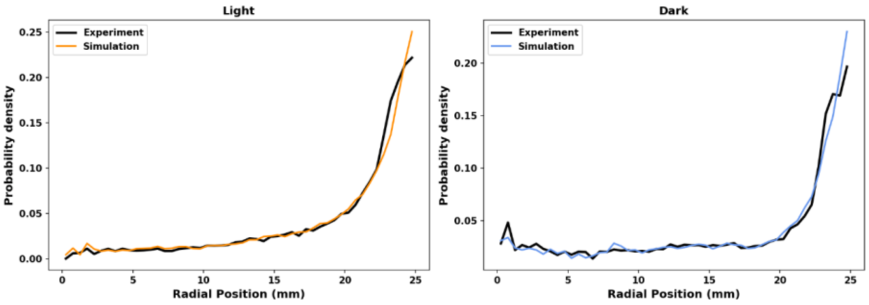


**Supplementary Figure 6.**
The probability distribution of radial position, p(*r*) for single fish in circular arenas of radius 25mm in (left panel) light and (right panel) dark, with experimental and simulation data plotted as in Supp. Fig. 3 and 4. The radial distribution function is area-normalized by 1/r.

### Simulating the motion of pairs of fish using single-fish properties plus pair properties derived from observations

As in the single fish simulations, the motion of each fish in the pair is modeled as a memoryless random walk with linear steps. The fish are treated as point objects; one can think of these as head positions and we will refer to the inter-fish distance as $d_{HH}$ (head-to-head distance) to match the terminology of the experimental data. As above, we make use of experimentally derived sets of bout and inter-bout parameters: bout duration (Δt), displacement magnitude or step size (Δs), and inter-bout intervals (T). These are binned by *r* and ψ, where r is the radial position and ψ is the wall alignment angle. Note that, as described below ("Kinematic parameters from pair data"), the final model jointly samples (Δs, Δt, T) from paired fish data, binned by (r, $d_{HH}$). Our use of turning angle (Δθ) is described two sections below.

**Commented [JE4]:** memoryless or memory-less? we have used both

**Commented [LD5]:** Maybe useful to specify which point is used on the body? I am assuming the head with the corrected calculation of its position? (I think this is explained in the methods and could be referred to here)

**Commented [RP6R5]:** Since they're points it doesn't correspond to anything; it's an abstraction of the fish. Still, the new text should clarify this.

### Simulating the motion of pairs of fish using single-fish properties only

Naively simulating the motion of two fish using two instances of the single-fish model unsurprisingly fails to reproduce the distribution of inter-fish head-to-head distances, p($d_{HH}$) (Supp. Fig. 7). These obviously non-interacting pseudo-pairs will also be useful as control sets, described below. The pseudo-pair p($d_{HH}$) has a strong peak at large separations, as expected given that the fish spend most of their time near the arena edge. (If one draws chords at random on the rim of a circle, the distribution of chord lengths is maximal, and in fact diverges, as the chord length approaches the circle diameter).

**Commented [LD7]:** Should we add a heading to title this section? for example 'Simulating the motion of pairs of fish using single-fish properties only'

**Commented [RP8R7]:** Done.

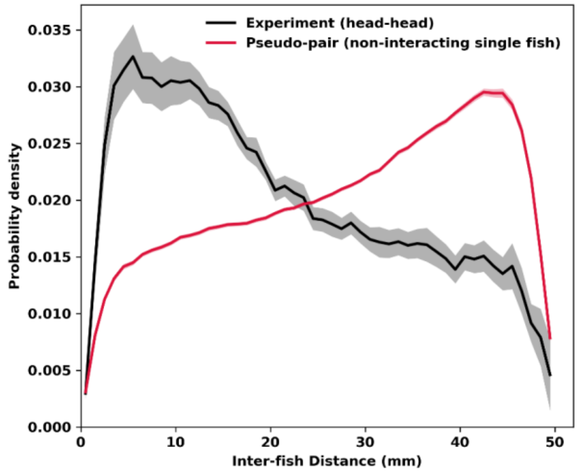


**Supplementary Figure 7.**
The experimental probability distribution of inter-fish distances, $p(d_{HH})$ (as in Supp. Fig. 3), together with the $p(d_{HH})$ constructed by taking pairs of single-fish datasets. As expected due to the absence of interactions in the latter, the two curves are very different.

Aspects of single-fish properties such as turning and kinematic behaviors must in reality be modulated in some way based on social parameters such as inter-fish distance ($d_{HH}$) and relative orientation ($\phi$). There are many ways to imagine this modulation can occur. Many possible models reproduce the observed $p(r)$. Some models can reproduce key features of the inter-fish distance distribution ($p(d_{HH})$), such as a region of highest probability from about 5 to 15 mm. Nearly all schemes we have tried, however, fail to reproduce *both* $p(r)$ and $p(d_{HH})$ unless various ad-hoc adjustable parameters are invoked. After careful assessment of how different models correspond to different assumptions about fish behavior and geometry, we converged on the model presented here, which gives very good agreement with the observed $p(r)$ and $p(d_{HH})$ distributions as well as the turning angle distribution, and which has **zero** relevant free parameters.

### Hypothesized social modulation of turning

A fish's turning angle in a simulated bout combines two components: an 'intrinsic' turning angle derived from the single fish data and a 'social' turning angle ascribed solely to the inter-fish geometry. The combination of the two angles involves a weighted sum with the weight determined from pair data and a modulation of the variance of the turns, also determined from pair data. We elaborate on all of these below.

**Commented [JE9]:** " or ' as used in other places in the MS

**Intrinsic turning angle.** The turning angles realized in experiments involving single fish are binned in r and ψ, giving at each (r, ψ) a set of empirical turning angles denoted $\{\Delta\theta_i\}$. At the current value of (r, ψ) in a simulated trajectory, we draw a turning angle $\Delta\theta_0$ at random from $\{\Delta\theta_i\}$. We also note $<\Delta\theta_i>$, the mean of $\{\Delta\theta_i\}$.

**Social turning angle.** The social turning angle $\Delta\theta_s$ at each step is defined as the angle by which the fish would have to turn so that the other fish would be directly in front of it; this is simply the relative orientation ($\phi$). It is conceivable, given the geometry of fish vision, that the preferred orientation is offset from directly forward; including such an offset of e.g. 20° or 45°, we find, has negligible impact on the results, likely because such an offset is necessarily symmetric about 0°.

**Weight.** The turning angle implemented at each step is a weighted sum of the intrinsic and social turning angles. The social weight (w) depends on the inter-fish distance ($d_{HH}$), and its magnitude is derived from the data itself. We tabulate from paired fish experiments the turning angles, $\Delta\theta_T$, i.e. the difference in angle between consecutive bout displacement (main text Fig**.** 3C), and bin by inter-fish distance and relative orientation ($d_{HH}$). We calculate w from fitting to the angles, in a rough sense considering $\Delta\theta_T = (1-w) <\Delta\theta_i> + w\ \Delta\theta_s$ and determining the weight that would best match the experimental observations. More precisely, this is done as a weighted least-squares projection of the observed per-bout turn vectors $z_{exp} = \exp(i\ \Delta\theta_T)$ onto the turn vectors $z_{int} = \exp(i <\Delta\theta_i>)$ and $z_{social} = \exp(i\ \Delta\theta_s)$. From

$$z_{exp} = (1-w)\, z_{int} + w\, z_{social}$$

i.e.,

$$z_{exp} - z_{int} = w\,(z_{social} - z_{int})$$

We pool the bout data into $d_{HH}$ bins and calculate the weight corresponding to each bin as the least-squares solution to the preceding equation:

$$w(d_{HH}) = \frac{\sum (z_{exp} - z_{int})(z_{social} - z_{int})^*}{\sum |z_{social} - z_{int}|^2}$$

where * indicates the complex conjugate.

**True Social Weight.** Importantly, the circular arena geometry itself is a major determinant of turning; if the arena edge is to the right of a fish, the other fish cannot be to the right, a turn cannot be to the right, and the correlation between social turning angle and relative orientation is not a true indicator of behavior. To identify and remove the geometric contributions to the turning angle weight, we make use of artificial pseudo-pair datasets constructed from two independent single-fish datasets, so that any correlations cannot possibly be due to real social interactions, while retaining the effects of arena geometry. The social weight derived from the pseudo-pair data is subtracted from the social weight derived from real experimental paired-fish data, keeping everything else unchanged. We denote the resulting “excess” weight, again a function of inter-fish distance, as $w_{excess}$:

$$w_{excess} = w_{pair} - w_{pseudopair}$$

We plot $w_{excess}$ vs. $d_{HH}$ for fish in light in Supp. Fig. 8 and for both light and dark in main text Fig. 3E. Note that in the light, this “genuine” social weight decays to zero by about 20 mm, while in the dark the social weight is shorter-ranged.

We use $w_{excess}$ to calculate a mean turning angle:

**Commented [LD10]:** Maybe should specify that this is the difference in angle between consecutive bouts, not within a bout. We should make that clear in the methods too. Do you think we should modify Fig. 1E accordingly? Are we using the bout turning angle (as opposed to bout-to-bout turning angle used for the maps in Fig. 3C) anywhere else?

$$\Delta\theta_M = (1 - w_{excess})\langle\Delta\theta_i\rangle + w_{excess}\Delta\theta_s$$

We explain the use of this $\Delta\theta_M$ in the next section, and alternative choices that do not give results that match the data later in this document.

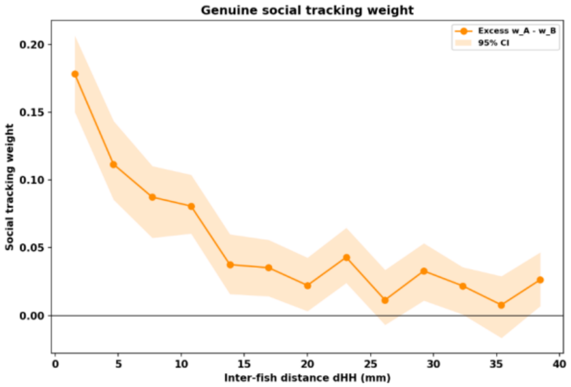


**Supplementary Figure 8.**
The genuine social weight $w_{excess}$ for pairs of fish in lit arenas, which decays from a maximum at very close separations to roughly zero by $d_{HH}$ = 20mm.

**Social Modulation of Turn Variance.** Surprisingly, we found that the variance of turning angles for fish in pairs shows a strong dependence on inter-fish distance and relative orientation, being broadest (high variance) at small distances and narrowest (low variance) at middle distances when the other fish is forward ($|\phi| < 90°$). We tabulated the experimental $\theta_T$ values in each ($d_{HH}$, $\phi$) bin and calculated the standard deviation; the resulting $std_\theta(d_{HH}, \phi)$ is plotted in Supp. Fig. 9 and main text Fig. 3D. We determined the large $d_{HH}$ asymptote, $std_{\theta,far}$ with an exponential fit of $std_\theta$ versus $d_{HH}$, pooled over $\phi$. To capture this modulation of turning variance in the simulation of pair behavior, we defined a focusing factor $f = std_\theta(d_{HH}, \phi) / std_{\theta,far}$ and calculated the turning angle used at each step of the paired fish simulations as

$$\Delta\theta = \Delta\theta_M + f\,(\Delta\theta_0 - \langle\Delta\theta_i\rangle),$$

where $\langle\Delta\theta_i\rangle$ is the mean of the intrinsic (single fish) turning angles, $\Delta\theta_i$.

A focusing factor $f = 0$ would correspond to zero angular spread, i.e. the fish moving in exactly the direction indicated by the weighted mean sum. A focusing factor $f = 1$ gives

$$\Delta\theta = \Delta\theta_0 + w_{excess}\,(\Delta\theta_s - \langle\Delta\theta_i\rangle) \quad [f = 1],$$

i.e., the turning angle is the value sampled from the single-fish distribution plus a bias towards the neighbor. Note that the use of the focusing factor $f$ preserves the functional form of the distribution of turns explored by a single fish, i.e. the set of possible $\Delta\theta$, while encoding the narrowing ($f < 1$) or widening ($f > 1$) of the distribution due to the observed influence of the neighbor. By sampling the empirical turn distribution, we make no assumptions about the form of the distribution (we find that the distribution is, in fact, non-Gaussian; see below).

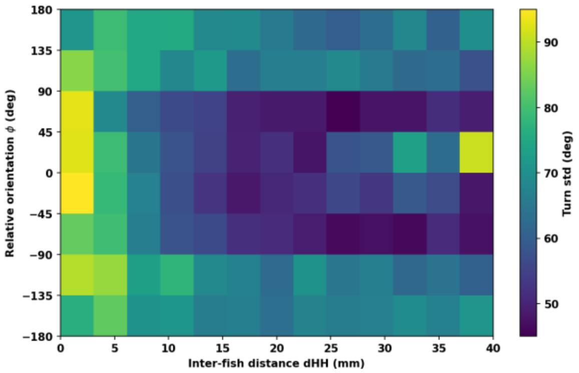


**Supplementary Figure 9.**
The standard deviation of the turning angle, $std_\theta(d_{HH}, \phi)$, in each ($d_{HH}$, $\phi$) bin. Note the low values at intermediate distances (roughly 10-25 mm) and orientations indicating a forward neighbor.

### Neglecting the social modulation of turning makes the model worse

Neglecting the socially induced modulation of turning leads to pair-distance histograms that are in poor accord with the experimental data.

**Removing the social weight.** Setting the weight $w_{excess}$ to zero (i.e., no social influence on the turn direction) but keeping the socially induced modulation of turning angle variance has a small effect on model performance; Supp. Fig. 10.

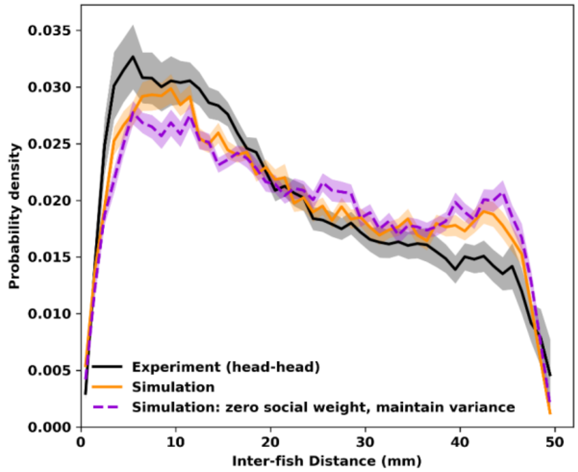


**Supplementary Figure 10.**
The experimental probability distribution of inter-fish distances, $p(d_{HH})$ (black), the results of the full model simulation (orange), both as in Supp. Fig. 3, together with the simulated $p(d_{HH})$ if the social weight $w_{excess}$ set to zero (i.e., removing any social influence on the turn direction) but retaining social influence on turn variance (violet).

**Removing the socially modulated variance.** Setting the focusing factor $f$ to 1 (i.e. assuming a constant variance in the fish turning angles, unaffected by the neighbor), has a dramatic effect on model agreement with the data; see Supp. Fig. 11.

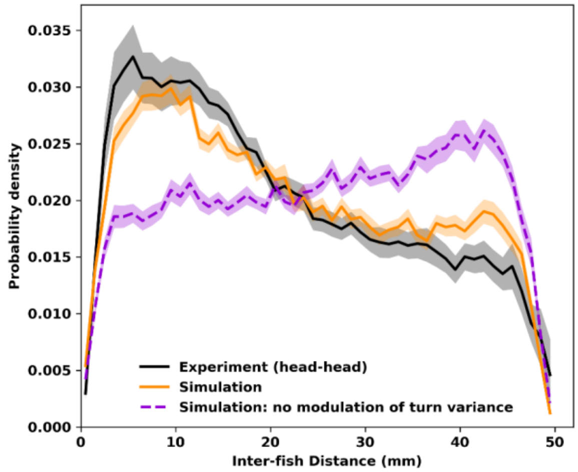

**Supplementary Figure 11.**
The experimental probability distribution of inter-fish distances, $p(d_{HH})$ (black), the results of the full model simulation (orange), both as in Supp. Fig. 3, together with the simulated $p(d_{HH})$ if the social modulation of turn variance $f$ is set to 1 (i.e., assuming a constant variance in the fish turning angles, unaffected by the neighbor; violet).

**Assuming Gaussian variance.** Using a Gaussian distribution of turning angles with standard deviation matched to the experimental observations at each $d_{HH}$ bin, i.e. keeping a socially modulated variance but replacing the true form of the variance with a Gaussian distribution, worsens the agreement between the model and the data; see Supp. Fig. 12.

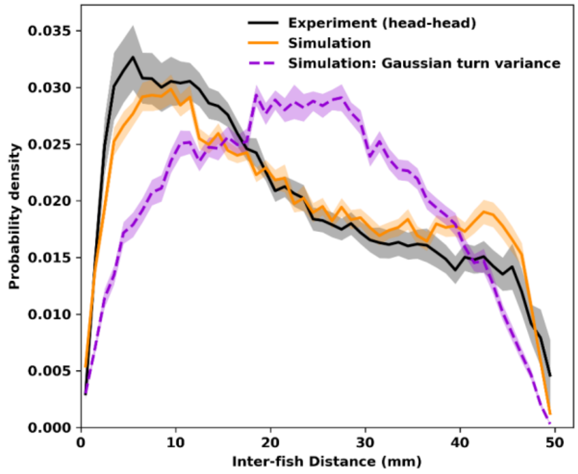


**Supplementary Figure 12.**
The experimental probability distribution of inter-fish distances, $p(d_{HH})$ (black), the results of the full model simulation (orange), both as in Supp. Fig. 3, together with the simulated $p(d_{HH})$ if the turn variance is modeled as Gaussian with the same standard deviation as the experimental data (violet).

**Using time-shifted rather than pseudo-pair data as the null for $w_{excess}$.** As described above, the true social weight $w_{excess}$ is obtained by subtracting from the apparent social weight the weight derived from control datasets constructed from pseudo-pairs of single fish datasets, obviously devoid of social interactions. Alternatively, one might use the control used in most of the main text which is constructed by taking real pair data and time-shifting the trajectory of one of the fish by half the movie duration. This is a flawed approach: the resulting $w_{excess}$ is smaller, and the resulting $p(d_{HH})$ has worse agreement with the data (Supp. Fig. 13). This is because the time-shifted pair data removes *instantaneous* real social interactions, but not the impact of social interactions accumulated over time (imagine, for example, that a result of instantaneous social interactions is fish moving more towards the center of the dish; on average, the time-shifted fish would also be distributed more towards the center, and calculating $w_{excess}$ using the time-shifted data would remove this true social effect). The time-shifted control is appropriate for its uses in the main text, of highlighting the role instantaneous social interaction, but it is inappropriate for calculating $w_{excess}$.

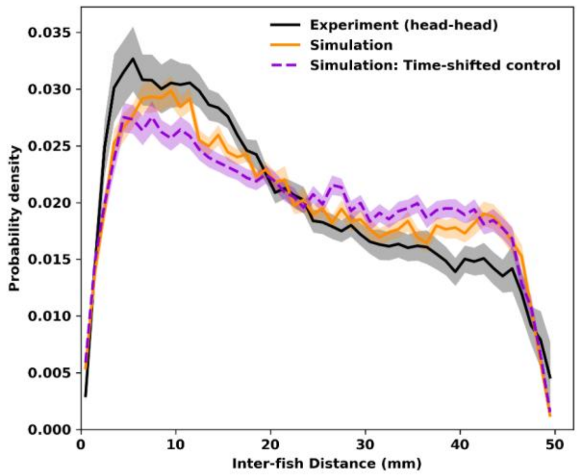


**Supplementary Figure 13.**

The experimental probability distribution of inter-fish distances, $p(d_{HH})$ (black), the results of the full model simulation (orange), both as in Supp. Fig. 3, together with the simulated $p(d_{HH})$ if time-shifted pair data, rather than pseudo-pair data, is used as the control.

### Kinematic parameters from pair data

We noticed that fish in pairs display different bout kinematic parameters, namely step sizes $\Delta s$, bout durations $\Delta t$, and inter-bout intervals T, than single fish and that the difference is inter-fish distance-dependent, especially manifesting as large motions when pairs are in close proximity (see main text Fig. 2, 4). We therefore jointly sample each of these kinematic parameters from the pair data, binned by (r, $d_{HH}$). Using instead kinematic parameters from the single-fish data has a negative impact on the correspondence between the simulation and the experimental data in pairs; see Supp. Fig. 14.

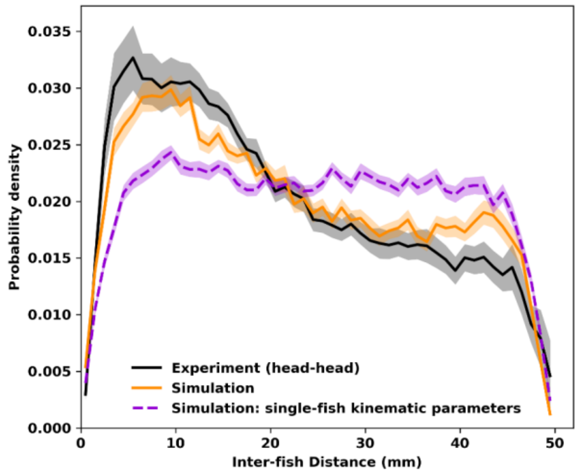


**Supplementary Figure 14.**
The experimental probability distribution of inter-fish distances, $p(d_{HH})$ (black), the results of the full model simulation (orange), both as in Supp. Fig. 3, together with the simulated $p(d_{HH})$ if single-fish kinematic parameters are used, rather than distance-modulated pair parameters.

### Accounting for tracking error; other parameters

The pair simulation model described above has zero free parameters, with bout and turning properties sampled from the data itself. Tracking errors lead to some unphysical steps. Incorporating conservative constraints on the resampling, excluding bouts with an apparent speed greater than 100 mm/s and, for the turning angle determination, steps of magnitude less than 1 mm, removes about 1% of the data and has a small but noticeable effect on $p(d_{HH})$; see e.g. Supp. Fig. 15, in which we don't exclude bouts with apparent speed greater than 100 mm/s.

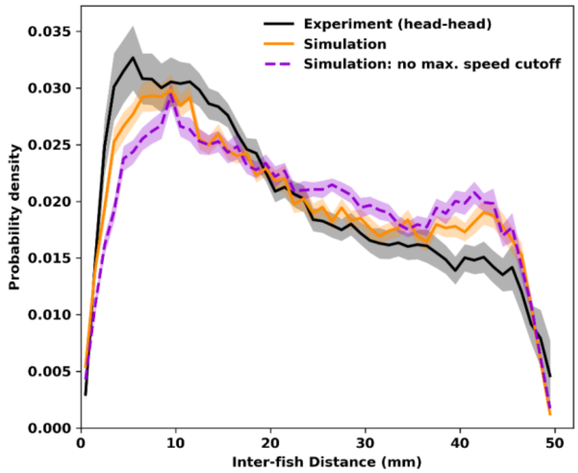


**Supplementary Figure 15.**
The experimental probability distribution of inter-fish distances, $p(d_{HH})$ (black), the results of the full model simulation (orange), both as in Supp. Fig. 3, together with the simulated $p(d_{HH})$ if we don't exclude bouts with apparent speed greater than 100 mm/s.

We also note that bin widths, the exponential form of the turning angle variance fit to $d_{HH}$, and the above-described tracking quality thresholds are modeling choices or parameters, but are not tuned to fit the target $p(d_{HH})$ and p(r) distributions.

### Additional Comments: Neglecting Memory

The model described above gives radial position distributions, p(*r*), and inter-fish distance distributions, $p(d_{HH})$, that are in very good, but not perfect, agreement with experimental observations for fish interacting in light (Supp. Fig. 3). Agreement in the dark is excellent. The success of the model in describing the data suggests that decision making by the two-week-old zebrafish is essentially instantaneous, not requiring memory or persistent behavioral states. Decisions in our model depend only on the current position and orientation of the fish relative to the arena and the other fish.

**Commented [JE11]:** elsewhere we say two-week-old

The remaining discrepancy between the model and the data, especially magnitude of the $p(d_{HH})$ peak being slightly lower around 5-15 mm in simulations than in reality, could be an indication of temporal correlations. Fish may execute persistent motions over multiple bouts, for example chasing each other at close separations, which would magnify $p(d_{HH})$ compared to the model. At large separations, i.e., on opposite sides of the dish, the model overstates the probability of fish maintaining large separations; this may also be due to a lack of persistence, as the simulated fish execute a relatively random back and forth walk along the arena edge. The lack of perfect correspondence between the model and the data is not surprising, and the remaining discrepancy may indicate future directions of study.
